# Loss of the adhesion protein Kindlin-1 stimulates tumor clearance via modulation of Tregs

**DOI:** 10.1101/2022.03.04.483014

**Authors:** Emily R Webb, Georgia Dodd, Esme Bullock, Morwenna Muir, Margaret C Frame, Alan Serrels, Valerie G Brunton

## Abstract

The adhesion protein Kindlin-1 is over-expressed in breast cancer where it has been shown to be associated with metastasis-free survival, however, the mechanisms involved are poorly understood. Here, we report that Kindlin-1 promotes anti-tumor immune evasion in a mouse model of breast cancer. Deletion of Kindlin-1 in Met-1 mammary tumor cells leads to tumor regression following injection into immunocompetent hosts. This was associated with a reduction in tumor infiltrating Tregs and impairment of their immune-suppressive activities in Kindlin-1 depleted tumors. Similar changes in T cell populations were seen following depletion of Kindlin-1 in the polyomavirus middle T antigen (PyV MT)-driven mouse model of mammary tumorigenesis. Analysis of cytokines secreted from the Met-1 cells identified a significant increase in IL-6 secretion when Kindlin-1 was depleted. Conditioned media from Kindlin-1 depleted cells lead to a decrease in the ability of Tregs to suppress the proliferation of CD8^+^ T cells, which was dependent on IL-6 and depletion of CD25^+^ Tregs resulted in a reduction of Met-1 tumor growth in mice. Overall, these data identify a novel function for Kindlin-1 in the regulation of anti-tumor immunity through cytokine regulation of Treg number and function.

## Introduction

Kindlin-1 is a four-point-one, ezrin, radixin, moesin (FERM) domain-containing adaptor protein that localises to focal adhesions, where it plays an important role in controlling integrin activation via binding to integrin β subunits ^1^. Loss-of-function mutations in the gene encoding Kindlin-1, *FERMT1*, leads to Kindler Syndrome, a rare autosomal recessive genodermatosis that causes skin atrophy, blistering, photosensitivity, hyper or hypopigmentation, increased light sensitivity and an enhanced risk of developing aggressive squamous cell carcinoma ^2,3^. However, Kindlin-1’s role in cancer is complex as it can also have a tumor-promoting role ^1,4^.

In breast cancer, Kindlin-1 expression is higher in tumor *versus* normal breast tissue and its expression is associated with metastasis-free survival ^5,6^. Consistent with its recognised role in regulating integrin-extracellular matrix interactions, Kindlin-1 controls both breast cancer cell adhesion, migration and invasion^5,7^. Kindlin-1 also regulates TGFβ signalling and epithelial to mesenchymal transition (EMT) in breast cancer ^6^. We have previously shown in the polyomavirus middle T antigen (PyV MT)-driven mouse model of mammary tumorigenesis, that loss of Kindlin-1 significantly delays tumor onset and reduces the incidence of lung metastasis ^7^. Mechanistically, Kindlin-1 stimulates metastatic growth in this model via integrin-dependent adhesion of circulating tumor cells to endothelial cells in the metastatic niche ^7^.

Kindlin-1 has also been shown to regulate inflammation in the skin of Kindler syndrome patients, where a number of pro-inflammatory cytokines are upregulated ^8,9^, and increased expression of genes associated with cytokine signalling have been reported ^10^. Progressive fibrosis of the dermis that follows inflammation in Kindler Syndrome is consistent with enhanced cytokine signalling, and the resulting ‘activation’ of fibroblasts leads to enhanced extracellular matrix deposition ^8,10^. Although inflammatory cytokines can play an important role in tumor progression, it is not known whether, and if so how, Kindlin-1 regulation of inflammatory cytokines in the tumor microenvironments influences tumor growth. Here we report that Kindlin-1 promotes an immunosuppressive and pro-tumorigenic microenvironment in a mouse model of breast cancer. Specifically, genetic deletion of the gene encoding Kindlin-1 leads to a reduction in tumor infiltrating Tregs and impairment of their immune-suppressive activities, in turn promoting tumor clearance and the generation of an immunological memory response. This implicates Kindlin-1 in a previously unrecognised function of immune-modulation in the breast cancer microenvironment *in vivo*.

## Results

### Loss of Kindlin-1 leads to tumor clearance and immunological memory

We used a syngeneic model in which *Fermt1* had been deleted in the Met-1 murine breast cancer cell line (Kin1-NULL) and to which either wild type Kindlin1 (Kin1-WT), or a mutant that is unable to bind β-integrin (Kin1-AA) were reintroduced ^7^. Tumor growth was monitored following subcutaneous injection of cells into both CD-1 nude immune-compromised and FVB (syngeneic) mice. Loss of Kindlin-1 led to reduced tumor growth in CD-1 nude mice (Figure 1A, B) with significant differences in tumor size noted from day 10 onwards. A similar reduced tumor growth rate in CD1 nude mice was seen following injection of human MDA-MB-231 cells in which Kindlin-1 was depleted using shRNA (Supplementary Figure 1). In contrast when Met-1 cells were injected into immune competent FVB mice, although Kindlin-1 loss led to a similar delay in tumor growth at day 10, there was complete tumor regression by day 19 (Figure 1C, D). Thus, Kindlin-1 is required for tumor growth of Met-1 cells in mice with a functional immune system, similar to what we reported previously for FAK-deficiency in a mouse model of squamous cell carcinoma (SCC) ^11^. Growth of the Kin1-AA mutant-expressing tumors was indistinguishable from Kin1-WT tumors in both CD1 nude and FVB mice (Figure 1A, C respectively), implying that integrin dependent functions of Kindlin-1 are not important for the growth of Met-1 tumors.

**Figure 1.**
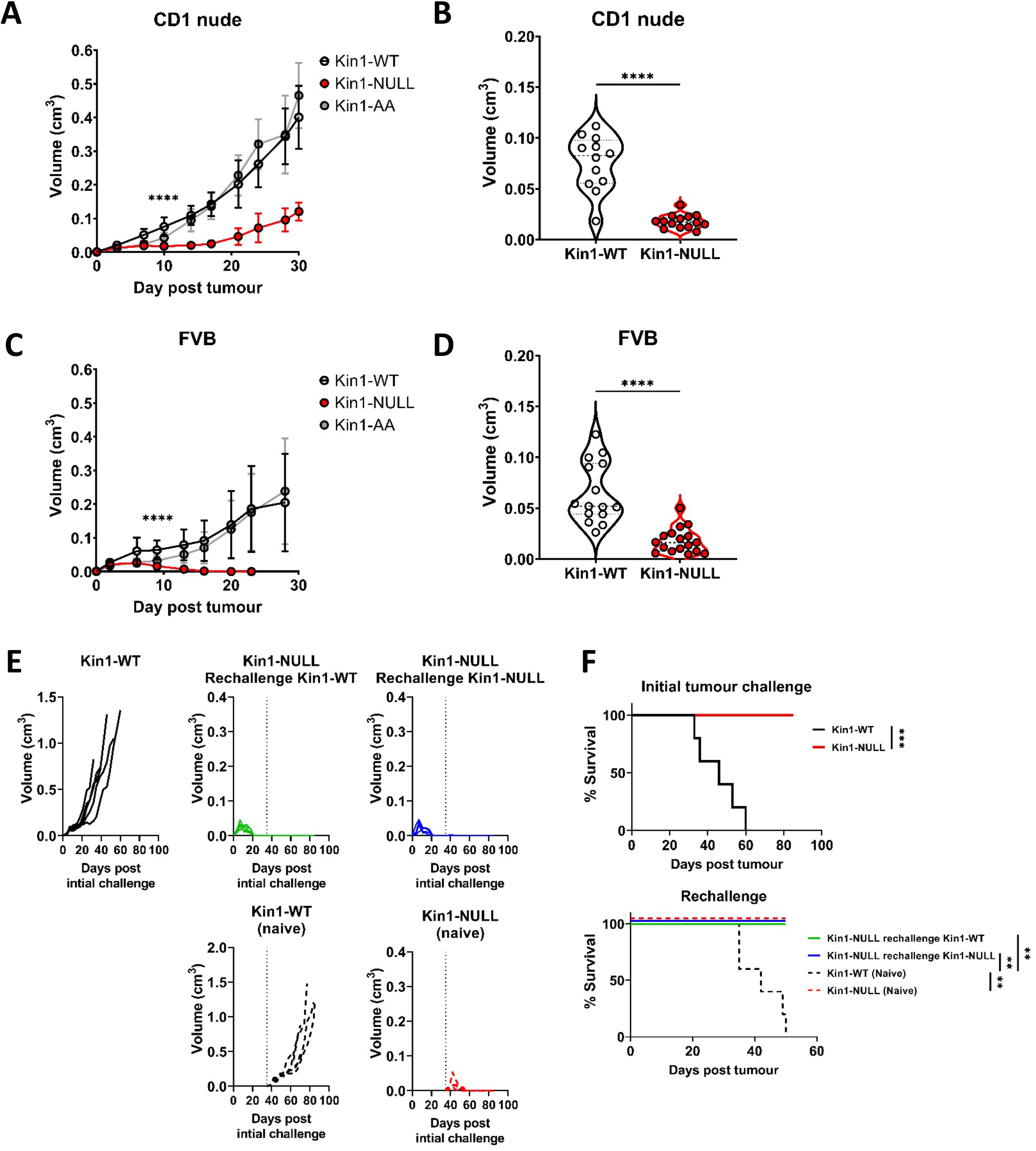
Loss of Kindlin-1 leads to tumor clearance and immunological memory. **A, C)** Met-1 Kin1-WT, Kin1-NULL or Kin1-AA tumors were established via subcutaneous injection into flanks of CD1 nude mice (**A**) or FVB mice (**C**). Tumor growth was monitored and recorded until day 30, with average tumor growth shown. **B, D)** Tumor size at day 10 post injection shown in CD1 nude mice (**B**) and FVB mice (**D**). **E)** Left flank of FVB mice was injected with Met-1 Kin1-WT or Kin1-NULL cells. At day 35 when no tumor was present, Kin1-NULL injected mice were rechallenged with either Kin1-WT or Kin1-NULL Met-1 cells on the right flank. Naïve FVB mice were also injected concurrently. Tumor growth and survival (**F**) were monitored throughout. Example of two independent experiments n=5-17 per group. Unpaired t-test (A-D) or Log Rank (F) with *= <0.05, ** = <0.01, *** = <0.001.

To further investigate whether an immune response was generated in mice with Kin1-NULL tumors, a re-challenge experiment was conducted. Following regression of Kin1-NULL tumors, mice were re-challenged with either Kin1-WT or Kin1-NULL cells on day 35 (Figure 1E, F). Neither Kin1-WT nor Kin1-NULL cells grew palpable tumors in mice which had been pre-challenged with Kin1-NULL cells, while injection of Kin1-WT or Kin1-NULL cells into naïve mice on the same day showed normal tumor growth and survival (Figure 1E, F). These data suggest that deletion of Kindlin-1 promotes effective immunosurveillance, resulting in tumor regression and lasting immunological memory. Furthermore, the inability of Kin1-WT cells to give rise to tumors in mice previously harbouring a Kin1-NULL tumor suggests that Kindlin-1 does not regulate key antigens permitting T-cell tumor recognition and effective immunosurveillance.

### Loss of Kindlin-1 modulates tumor associated myeloid populations

To understand how loss of Kindlin-1 promotes immunosurveillance, flow cytometry was used to profile the immune landscape of Kin1-WT and Kin1-NULL tumors at 10 days post tumor challenge. The percentages of major myeloid subsets were quantified within the tumors (Figure 2A, Supplementary Figure 2). Of note there were significantly reduced CD45^+^ cells within Kin1-NULL tumors, alongside a reduction in both monocytes and macrophages (Figure 2A). However, no significant differences were observed in expression of the phenotype markers MHC II, CD206 and SIRPα, between Kin1-WT and Kin1-NULL tumor-associated macrophages (Supplementary Figure 3A), suggesting that there is no change in the ‘polarisation’ status of these cells. Although there was no difference in total dendritic cell (DC) percentages, analysis of DC subsets demonstrated a significant increase in conventional type I DCs (cDC1) within Kin1-NULL tumors compared to Kin1-WT (Figure 2B, C). cDC1s are efficient at cross presentation, essential for CD8^+^ responses and have been demonstrated to be important for anti-tumor immune responses ^12,13^. Furthermore, we observed increased expression of the T-cell co-stimulatory molecule CD80 on DCs in Kin1-NULL tumors (Figure 2D), and increased expression of the T cell inhibitory PD-1 receptor ligand PD-L1 (Figure 2E). Additionally, analysis of bulk tumor RNA demonstrated an increase in antigen presentation (*H20Q2* and *H2-Eb1*) and antigen transport *(Tap2)* related genes within Kin1-NULL tumors (Figure 2F). An increase in *Ifng* was also seen in the Kin1-NULL tumors (Figure 2G), consistent with increased expression of IFNγ-inducible PD-L1, and MHC/Antigen processing genes ^14,15^. These data suggest that loss of Kindlin-1 may result in increased cross-presentation of tumor antigen by DCs, promoting T-cell activation and antitumor immunity. Despite an increase in PD-L1 protein expression noted on cDC1 cells, overall PD-L1 gene *(Cd274)* expression was found to be decreased on CD45^+^ cells isolated from both tumors and draining lymph nodes (dLN) of Kin1-NULL tumors, compared to Kin1-WT tumors (Figure 2H), and treatment of Kin1-WT tumor bearing mice with a PD-L1 blocking antibody, lead to a significant reduction in tumor size (Figure 2I). Analysis of a publicly available human breast cancer data set (METABRIC), demonstrated a small but significant correlation between *FERMT1* (Kindlin-1) and *CD274* (PD-L1) gene expression (Supplementary Figure 3B). These data show that loss of Kindlin-1 leads to modulation of PD-L1 expression on tumor infiltrating immune cells, and that the PD-1/L1 pathway is important for Kin1-WT tumors to control the anti-tumor immune response.

**Figure 2.**
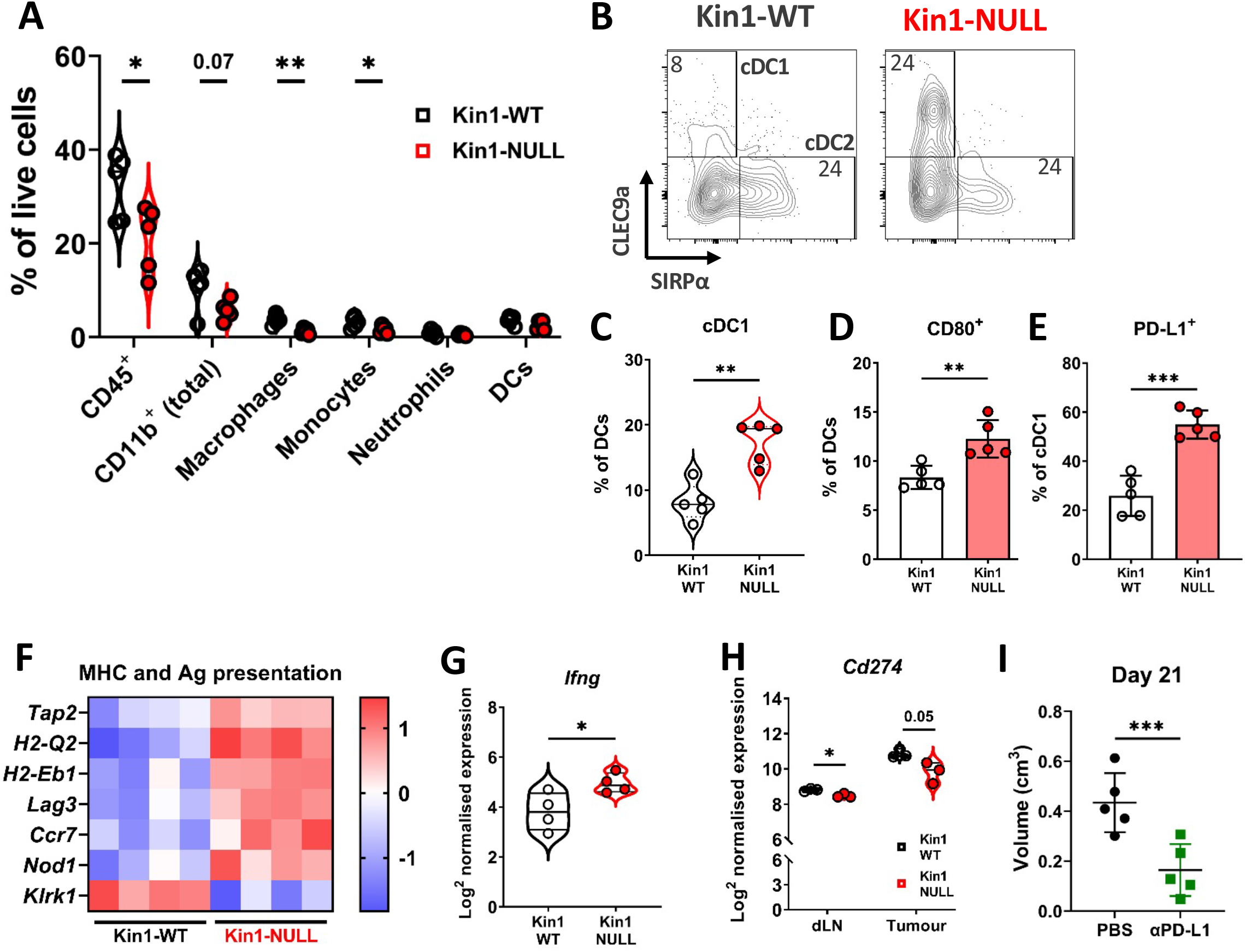
Loss of Kindlin-1 reduces tumor associated macrophages and increases cDC1 dendritic cells. **A)** Met-1 Kin1-WT or Kin1-NULL tumors were established via subcutaneous injection in FVB mice, and harvested at day 10 for immunophenotyping by flow cytometry. Major myeloid populations were quantified as a percentage of live (total) cells. **B)** Raw FACS plots demonstrating gating of cDC1 and cDC2 cells, and quantified **(C)** as a percentage of total DCs (CD11c^+^ MHC II^+^). **D)** Quantification of CD80 expression on total DCs by flow cytometry. **E)** Quantification of PD-L1 expression on cDC1 cells. **F)** As in A but bulk tumors were harvested for RNA expression analysis using Nanostring PanCancer Immune panel. Differentially expressed genes related to the gene sets ‘MHC’ and ‘Antigen (Ag) presentation’ are shown. **G)** Expression of *Ifng* using Nanostring PanCancer Immune panel comparing MET-1 Kin1-WT and NULL cells. **H)** Expression of *Cd274* (PD-L1) on isolated CD45+ cells using Nanostring Immune Exhaustion panel comparing draining lymph nodes (dLN) and tumors from MET-1 Kin1-WT and NULL tumor bearing mice. **I)** Tumors were established as in A, with mice concurrently treated with anti-PD-L1 antibody. Tumor size at Day 21 post tumor is shown. Example of two independent experiments (A-D+F), n=3-5 per group. For E, fold change cut off = 1.2, FDR = <0.05. Unpaired t-test with *= <0.05, ** = <0.01, *** = <0.001.

### Loss of Kindlin-1 reduces Treg infiltration and memory phenotype

The modulation of PD-L1 in Kin1-NULL tumors, and generation of immunological memory, suggested that the tumor clearance may be tied to a T cell directed anti-tumor immune response. To that end, further immunophenotyping of Kin1-WT and Kin1-NULL tumors was conducted with focus on T cell subsets (Supplementary Figure 4). Tumors were taken at day 10 post tumor cell implantation. We found a significant reduction of total CD3^+^ cells in Kin1-NULL tumors, which was driven by a decrease in CD4^+^ T cells (Supplementary Figure 5A). Within the CD4^+^ cell compartment, a significant decrease of regulatory T (Treg) cells as a percentage of total cells was evident in Kin1-NULL tumors compared to Kin1-WT tumors (Figure 3A). The reduction of Tregs was also observed when analysing cells as a percentage of total CD3^+^ cells, with a significant increase in non-Treg CD4^+^ cells as a proportion of total CD3^+^ (Supplementary Figure 5B). As Treg cells are widely reported to control anti-tumor T cell responses ^16^, these data suggest that Kindlin-1 loss results in fewer infiltrating suppressive T cells within the tumor microenvironment. Furthermore, the reduction in Tregs was mirrored in the tumor draining lymph nodes of Kin1-NULL tumors at day 10, but not observed systemically in the spleen (Supplementary Figure 5C). This suggests that the reduction in Tregs is due to changes in the local tumor environment.

**Figure 3.**
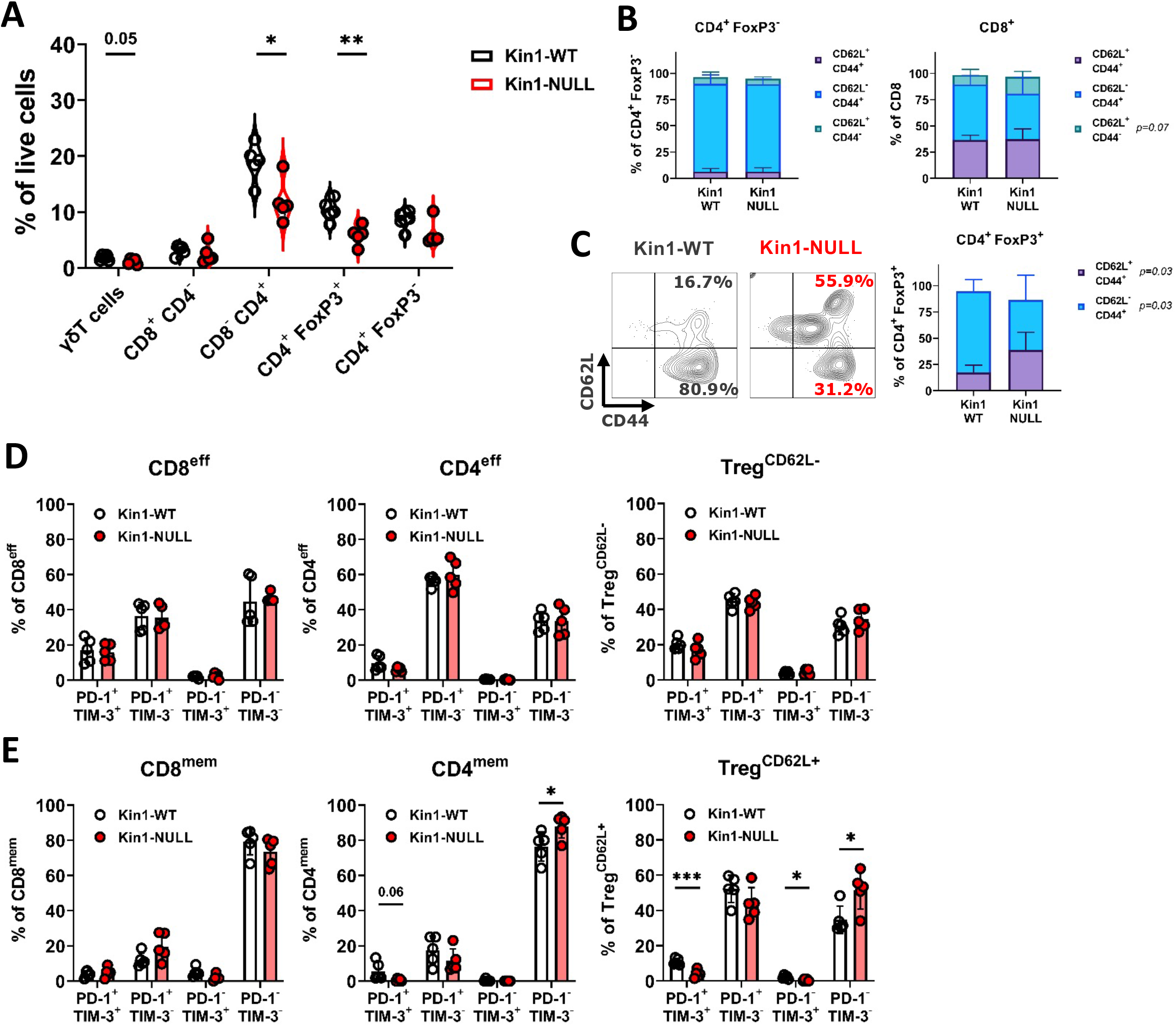
Loss of Kindlin-1 reduces tumor infiltrating Treg cells. **A)** Met-1 Kin1-WT or Kin1-NULL tumors were established via subcutaneous injection in FVB mice, and harvested at day 10 for immunophenotyping by flow cytometry. Gating of major T cell populations was conducted and quantified as percentage of total (alive) cells. **B)** Quantification of effector (CD62L-CD44+), memory (CD62L+ CD44+) and naïve (CD62l+ CD44-) populations as a percentage of corresponding T cell subset. **C)** Representative example of gating resting Tregs (CD62L+) and activated Tregs (CD62L-) in tumors, with quantification on the right. **D, E)** Quantification of PD-1 and TIM-3 expression on T cell subset effector (or CD62L-) populations (**D**) and memory (or CD62L+) populations (**E**). Example of two independent experiments (A-D). n=3-5 per group. Unpaired t-test with *= <0.05, ** = <0.01, *** = <0.001.

To further elucidate whether any T cell phenotypic changes occurred upon loss of Kindlin-1 in tumors, analysis of memory markers (Figure 3B, C) and activation/exhaustion markers (Figure 3D, E) was conducted. We found no significant changes in memory marker expression on non-Treg CD4^+^ cells; however, an increase in naïve (CD62L^+^ CD44^-^) CD8^+^ cells was observed (Figure 3B). Interestingly, a significant increase in CD62L^+^ Treg cells were evident within Kin1-NULL tumors when compared to Kin1-WT tumors at day 10 (Figure 3C). A corresponding reduction in CD62L^-^ Treg proportions were also noted. CD62L^+^ Treg cells have been categorised into resting Tregs, with reduced proliferative capacity, whereas CD62L^-^ populations are reported to be activated Tregs with increased suppressive capacity, and accumulation of these cells into tumors drive CD8^+^ T cell suppression ^17–19^. The switch in the prominence of these two subsets in the Kin1-NULL tumors suggests that more resting Tregs are present compared to Kin1-WT tumors. We found minimal changes in activation (PD-1+ TIM-3^-^) and exhaustion (PD-1+ TIM-3+) markers on effector (or CD62L-Tregs) (Figure 3D) non-Treg T cells. Of note, a reduction of double positive (exhaustion) cells within CD4^mem^ and Treg^CD62L+^ populations was seen in Kin1-NULL tumors, alongside a corresponding increase in double negative (PD-1^-^ TIM-3^-^) cells (Figure 3E). These data suggest that loss of Kindlin-1 in Met-1 tumors is primarily modulating CD4^+^ T cell phenotypes, specifically that of Tregs.

### Loss of Kindlin-1 also causes immune changes in a spontaneous breast cancer model

To investigate whether similar immune modulation is evident when Kindlin-1 is depleted in a spontaneous mammary tumor model, immunophenotyping of tumors from the MMTV-PyV MT mouse model was carried out. MMTV-PyV MT mice were crossed with mice in which exons 4 and 5 of the *Fermt1* gene were flanked with LoxP1 recombination sites, and in which Cre recombinase was expressed in the mammary epithelium under transcriptional control of the mouse mammary tumor virus (MMTV) ^7^. Tumors that formed were collected from MMTV-PyV MMTV-Kin-1^wt/wt^ (MT-Kin-1^wt/wt^) and MMTV-PyV MMTV-Kin-1^fl/fl^ (MT-Kin-1^fl/fl^) mice and immune cell populations analyzed. We observed an increase of cDC1 cells in tumors from the MT-Kin-1^fl/fl^ mice when compared to tumors in the MT-Kin-1^wt/wt^ mice (Supplementary Figure 6A), similar to the Met-1 cell line-derived tumors already described (Figure 2B). Furthermore, there was a reduction of PD-L1 expression on CD45^+^ cells in MT-Kin-1^fl/fl^ tumors (Supplementary Figure 6B), as well as reduced PD-L1 on several myeloid subsets, demonstrating modulation of the PD-L1 pathway in Kindlin-1 deficient tumors also in this spontaneous breast cancer model.

Analysis of T cell subsets demonstrated significant reduction of T cells in MT-Kin-1^fl/fl^ tumors as a percentage of total cells when compared to MT-Kin-1^wt/wt^ (Supplementary Figure 6C), although there was large variability between animals. This may be due to the asynchronous growth characteristics of the spontaneous MMTV model, with some of the mammary tumors enveloping nearby lymph nodes as they progress. Of note, total CD3 infiltration of these tumors was lower than in the Met-1 model (Supplementary Figure 5A). However, when analyzed as a percentage of total CD3^+^ cells, MT-Kin-1^fl/fl^ tumors had significantly fewer Treg cells with an increase in percentage of non-Treg CD4+ cells (Supplementary Figure 6D), that was similar to the Met-1 cell line derived tumor model (Supplementary Figure 5B). These data demonstrate immune modulation upon the loss of Kindlin-1 tumor expression in distinct models of breast cancer, with consistent changes in Treg and cDC1 cells resulting from Kindlin deficiency. However, in the MT-Kin-1^fl/fl^ mice we do not see tumor regression, which most likely reflects the incomplete loss of Kindlin-1 in the mammary tumor cells in this model ^7^.

### Kindlin-1 knock-out cells modulate Treg phenotype and function

As Tregs, which drive suppression of effector T cell function, were observed to be reduced in number in Kin1-NULL tumors, assessment of immunosuppressive pathways was conducted. Analysis of RNA from CD45^+^ tumor infiltrating cells isolated from Kin1-WT and Kin1-NULL tumors was carried out which showed a reduction of various T cell inhibitory checkpoint pathway related genes, including *Pdl1, Vista and Tim3* in Kin1-NULL tumors (Figure 4A). As many of these suppressive receptor pathways are known to be utilised by Treg cells for effector cell suppression, we next addressed whether Kindlin-1 deficiency leads to widespread disruption of Treg phenotype and function. Detailed Treg profiling from the Met-1 cell line derived tumors at day 10 by flow cytometry, was carried out (Figure 4B-D and Supplementary Figure 7). Of note, downregulation of TNF superfamily co-stimulatory receptors GITR, 4-1BB and OX40 was observed in Kin1-NULL infiltrating Tregs (Figure 4B), which are critical for Treg development and promoting their proliferation ^20,21^. Expression of the inhibitory receptors LAG-3, CTLA-4, TIGIT and PD-1 were also downregulated on the surface of Tregs in Kin1-NULL tumors (Figure 4C). Although ligation of these receptors in CD8^+^ and non-Treg CD4^+^ cells inhibits effector function, these receptors are crucial for Treg differentiation and immunosuppressive activity ^21^. There was significant downregulation of both CD73 and CD39 expression on Tregs from Kin1-NULL tumors (Figure 4D). The CD39/CD73 pathway is a major modulator of Treg activity via metabolism of ATP to create extracellular adenosine, in turn inhibiting effector T cell function ^22,23^. This suggests that loss of Kindlin-1 may cause metabolic changes in Treg cells, resulting in impairment of their suppressive capacity. Taken together, down regulation of these phenotypic markers suggests that Tregs from Kin-1 NULL tumors could be less immunosuppressive, and therefore allow development of a sufficient anti-tumor immune response, leading to tumor clearance. To that end, analysis of activation markers on CD8+ T cells was assessed. Although, we have previously shown little modulation of CD8^+^ cell number and expression of PD-1 and TIM-3 (Figure 2), this expanded panel allowed for assessment of CD8 activation in greater depth. There was an increase in expression of OX40, CD83 and CD29 on CD8^+^ T cells infiltrating Kin1-NULL tumors compared to those in Kin1-WT tumors (Figure 4E). These receptors are associated with activated T cells and an increase in cytotoxic potential ^20,24,25^ and these data suggest that loss of Kindlin-1 causes a reduction in Treg suppressive function, which may enhance the activation of CD8+ cytotoxic T cells, leading to reduced tumor growth.

**Figure 4.**
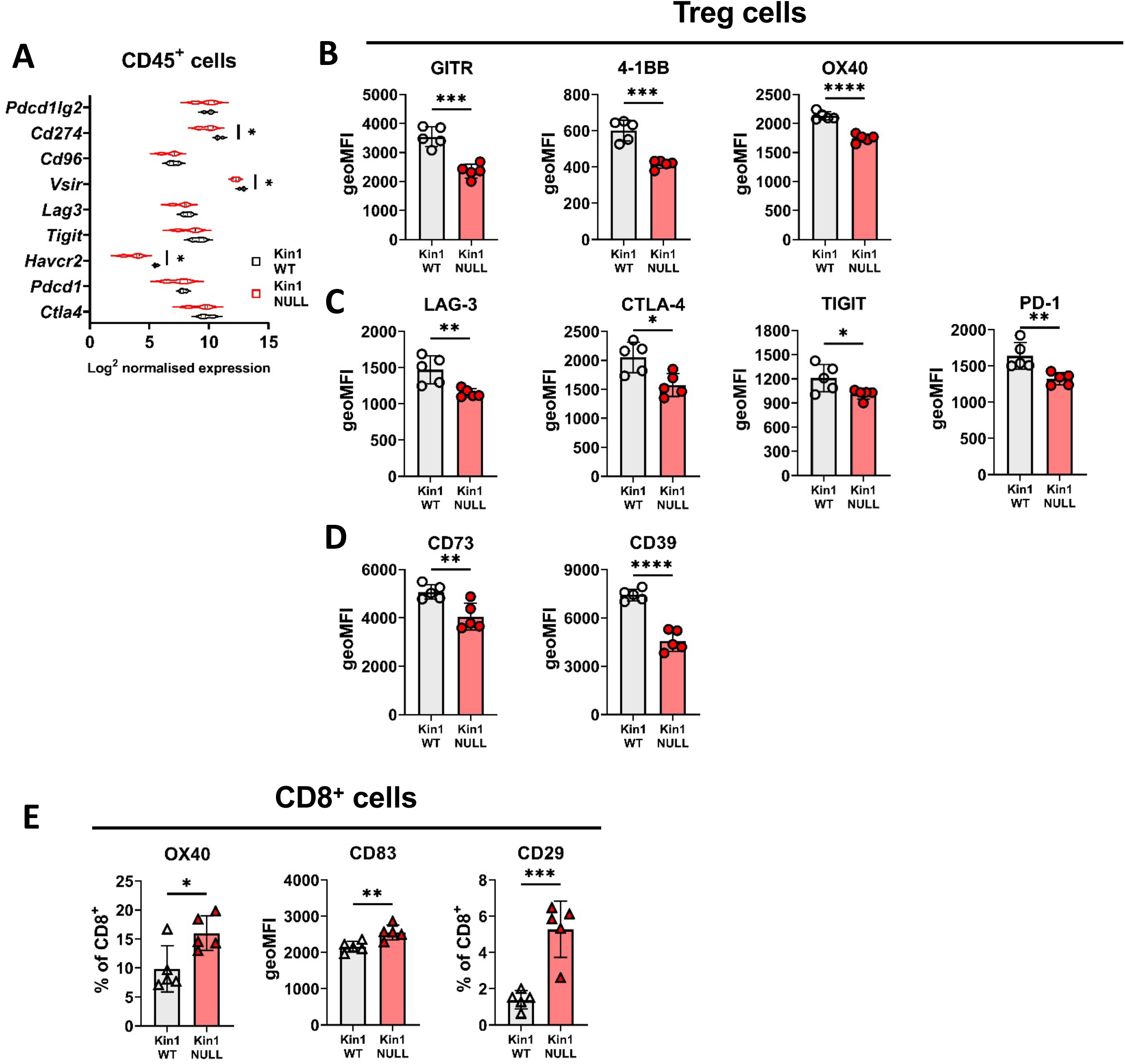
Loss of Kindlin-1 modulates Treg phenotype and function. **A)** Met-1 Kin1-WT or Kin1-NULL tumors were established via subcutaneous injection in FVB mice, and harvested at day 10 for RNA analysis using Nanostring Immune Exhaustion panel. Shown is log2 normalised expression of known T cell inhibitory receptors and pathways. **B)** As in A but tumors harvested for immunophenotyping by flow cytometry. Analysis of expression of markers were assessed on gated CD4+ FoxP3+ T cells (Tregs). Quantification of expression as geo mean fluorescent intensity (geoMFI) shown for TNF superfamily members (B), known inhibitory receptors **(C)** and metabolism related receptors **(D)**. Percentage expression is shown in Supplementary Figure 6. **E)** As in A with quantification of activation associated receptors on tumor infiltrating CD8^+^ T cells n=4-5 per group. Unpaired t-test with *= <0.05, ** = <0.01, *** = <0.001, **** = <0.0001.

### Loss of Kindlin-1 leads to altered cytokine secretion and regulates Treg differentiation

To understand the mechanism by which Kindlin-1 can regulate immune cell populations, we examined conditioned media collected from Kin1-WT and Kin1-NULL Met-1 cells using a forward phase protein array of 64 cytokines. Decreases in CXCL11, 12 and IL-12 and significant increases in CXCL13, IL-1RA and IL-6 were detected in conditioned media from Kin1-NULL cells compared to Kin1-WT cells (Figure 5A). Although CXCL13, a chemokine involved in B cell migration ^26^, was greatly increased, no significant increase in B cells within Kin-1 NULL tumors was observed (Supplementary Figure 8). We therefore focussed on IL-6 as previous studies have demonstrated the importance of IL-6 in influencing the differentiation of naïve CD4^+^ T cells into Tregs ^27^ and Treg suppressive function in various settings. ^28–31^ The increase in secretion of IL-6 protein in conditioned media from Kin1-NULL cells was confirmed by ELISA (Figure 5B), while bulk tumor RNA analysis of Kin1-WT and Kin1-NULL tumors confirmed changes in IL-6-related genes, with an increase in *Il6, Il6ra* and *Il6st* found in Kin1-NULL tumors (Figure 5C). Overall, this suggests that the altered cytokine profile secreted by Kin1-NULL cells is able to modulate signalling within local immune microenvironments.

**Figure 5.**
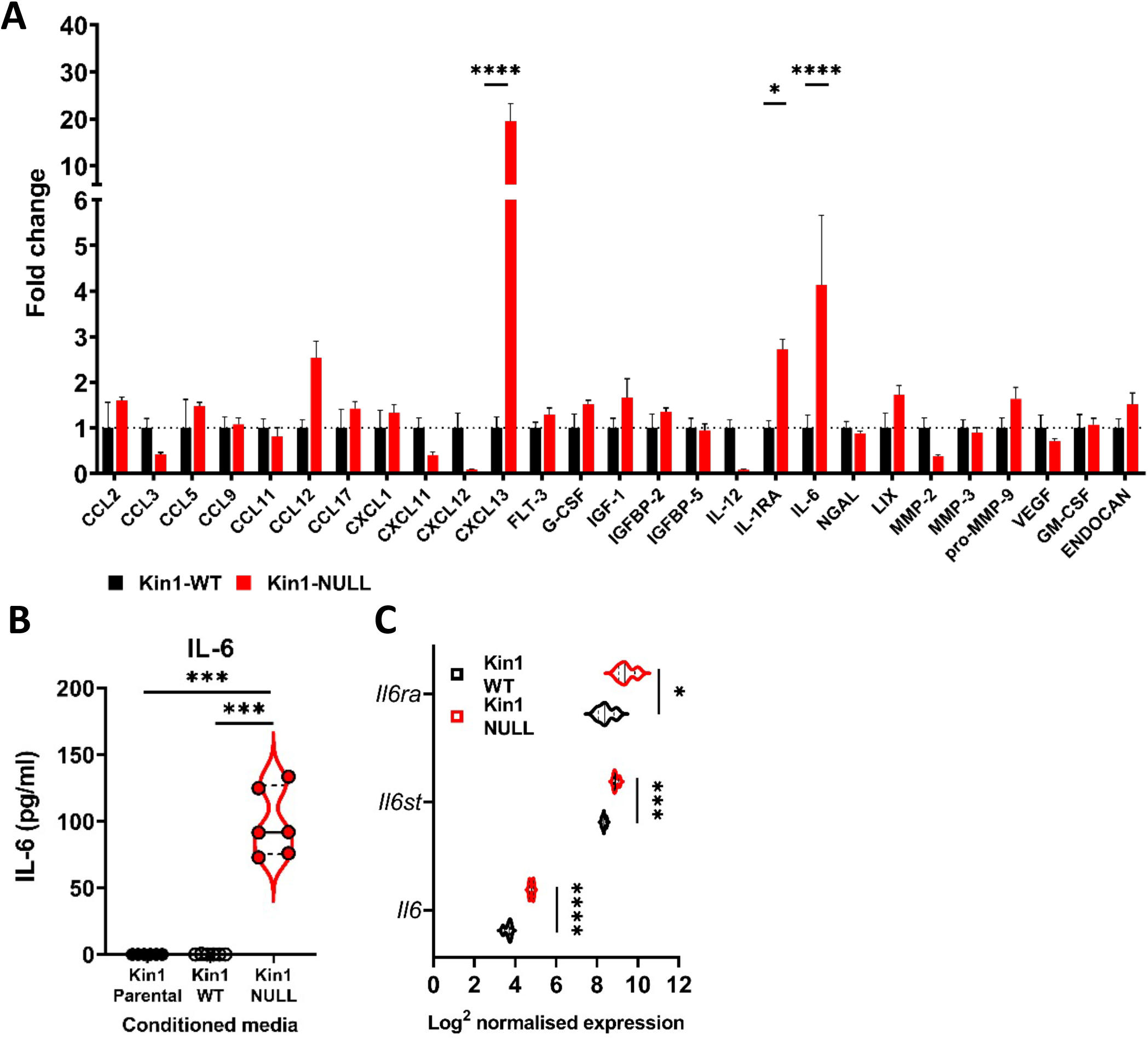
Loss of Kindlin-1 leads to altered cytokine secretion. **A)** Met-1 Kin1-WT or Kin1-NULL cells were cultured for 48hrs before conditioned media (CM) was harvested for analysis by forward phase protein array. Proteins detected above background are shown as fold change over Kin1-WT. **B)** Bulk tumour RNA analysis of Met-1 Kin1-WT or Kin1-NULL tumours at day 10. Log2 normalised expression of IL-6 related genes are shown. **C)** Quantification of IL-6 in Met-1 conditioned media via ELISA. Unpaired t-test with *= <0.05, ** = <0.01, *** = <0.001, **** = <0.0001.

Previous studies have demonstrated the importance of IL-6 in influencing the differentiation of naïve CD4^+^ T cells into either Tregs or Th17 cells when in the presence of TGFß ^27^. We therefore addressed whether tumor cell conditioned media could influence the differentiation of naïve CD4^+^ T cells *in vitro*. Conditioned media from Kin1-NULL cells resulted in a reduction of differentiation of CD4^+^ T cells into FoxP3^+^ Tregs compared to that seen following incubation with conditioned media from Kin1-WT cells (Figure 6A, B). The decrease in CD4^+^FoxP3^+^ cells was accompanied by a corresponding increase in CD4^+^RoRγT^+^ Th17 cells (Figure 6C), implying that the conditioned media from Kin1-NULL cells is diverting naïve CD4^+^ differentiation towards a more Th17 cell than Treg cell phenotype. When we looked at RNA expression in CD45^+^ cells isolated from Kin1-NULL and Kin1-WT tumors, we found a significant increase in Th17 associated gene, *Rorg* ^32^, in Kin1-NULL tumors compared to Kin1-WT (Figure 6D) supporting a role for Kindlin-1 in modulating CD4+ T cell differentiation.

**Figure 6.**
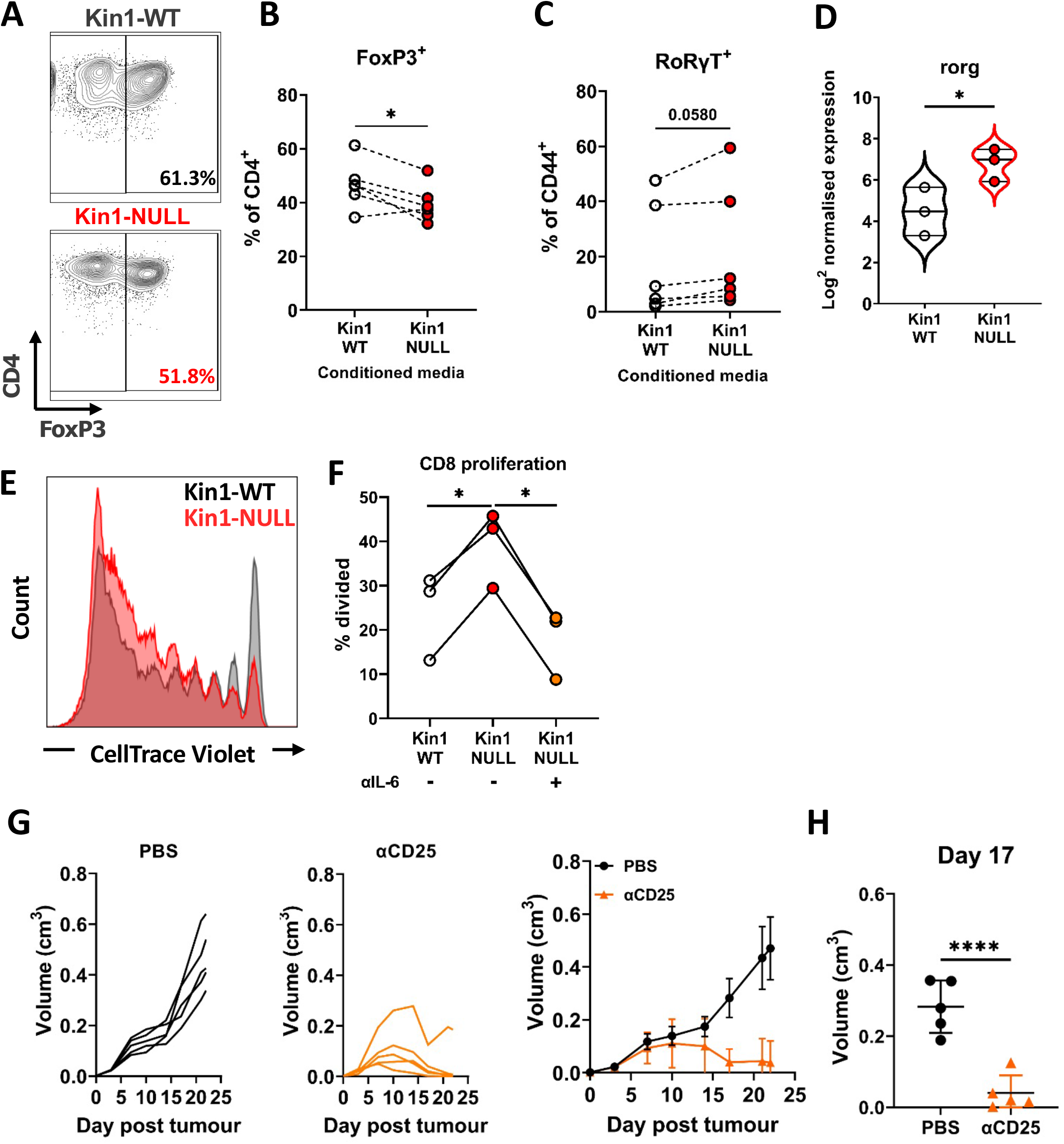
Loss of Kindlin-1 leads to modulation of Treg differentiation and function. **A-C)** Met-1 Kin1-WT or Kin1-NULL cells were cultured for 48hrs before conditioned media (CM) was harvested. Naïve CD4+ T cells were isolated from FVB mice spleens and stimulated in the presence of either Met-1 Kin1-WT or Kin1-NULL CM. At day 5 T cells were harvested for analysis of Treg differentiation by expression of FoxP3 (**A**). Example of gating is shown together with, **B)** quantification as percentage of CD4+ cells. Quantification of RoRγT expression as a percentage of CD4+ FoxP3-CD44+ cells (**C**). **D)** *Rorg* gene expression in isolated CD45+ cells from either Met-1 Kin1-WT or Kin1-NULL tumors as shown as Log2 normalised expression. **E)** CD8+ CD4-CD25- and CD8-CD4+ CD25hi (Treg) cells were sorted from FVB spleens. CD8+ cells were labelled with CellTrace Violet and co-cultured with Tregs under stimulation at a ratio of 1:8 (Treg:CD8), in the presence of conditioned media +/- anti-IL-6 blocking antibody. At day 5 cells were harvested and analysis of proliferation of CD8 cells was conducted. Example histogram of CellTrace Violet staining with **F)** quantification of CD8 proliferation shown. **G)** Met-1 tumors were established in mice pre-treated with anti-CD25 to deplete Tregs. Tumor growth for individual mice (left+middle) and averages (right) are shown. **H)** Tumor size at Day 17 from G. n=3-7 per group. For D, fold change cut off = 1.2, FDR = <0.05. Unpaired t-test with *= <0.05, ** = <0.01, *** = <0.001, **** = <0.0001.

To establish whether proteins secreted by Kin1-NULL cells can impair the ability of Tregs to suppress the proliferation of CD8+ T cells, a Treg suppression assay was conducted with addition of conditioned medium from either Kin1-WT or Kin1-NULL cells. Conditioned media from Kin1-NULL cells lead to a decrease in Treg suppressive capacity, shown as an increase in percentage of divided CD8^+^ T cells compared to incubation with Kin1-WT conditioned media (Figure 6E, F). Treatment with an antibody that blocks the function of IL-6 reduced the ability of conditioned media from Kin1-NULL to increase division of CD8^+^ T cells. Thus, loss of Kindlin1 in the Met-1 cells leads to increased secretion of IL-6, which in turn reduces the ability of Tregs to suppress CD8^+^ T cell proliferation. Finally, to demonstrate the importance of Tregs in controlling the anti-tumor immune response in Kin1-WT tumors, mice were depleted of CD25^+^ cells before tumor implantation (Supplementary Figure 9). Here, depletion of CD25^+^ cells resulted in a reduction of tumor growth compared to controls (Figure 6G), demonstrating a similar growth pattern as seen in Kin-1 NULL tumors (Figure 1C). There was a significant reduction in tumor size at Day 17 (Figure 6H) in Kin1-WT tumors with depleted Tregs. Overall, these data demonstrate the importance of Treg-mediated immune suppression in Kin1-WT tumors, leading to an increase in tumor growth. Deletion of Kindlin-1 from these tumors reduced both Treg number and function, and resulted in an increase in CD8^+^ activity and tumor clearance.

## Discussion

Previous studies have identified an important pro-tumorigenic role for Kindlin-1 in breast cancer where it promotes cell, migration, adhesion and EMT, and is associated with increased pulmonary metastasis and lung metastasis-free survival^5–7^. Here we show that Kindlin-1 can also regulate breast cancer progression by modulating the anti-tumor immune response through regulation of the immune composition of the tumor microenvironment.

Kindlin-1 is part of a family of proteins consisting of Kindlin-1, −2 and −3. They bind β integrin subunits and are required for integrin activation ^1^, and most studies have reported on their integrin-dependent roles in cellular phenotypes such as cell adhesion, migration and invasion. However, integrin-independent functions of Kindlin-1 have also been reported, where it can initiate downstream signalling out with integrin adhesions ^33,34^. Here we used a mutant of Kindlin-1 (Kin1-AA) that cannot bind β integrins, which we have previously shown leads to reduced levels of activated β1 integrin and integrin-dependent adhesion in Met-1 cells ^7^, and show that the ability of Kindlin-1 to bind integrins is not required for the growth of Met-1 tumors, or their immune clearance in immunocompetent animals. Redundancy between Kindlin-1 and Kindlin-2 has been reported in relation to integrin-dependent functions, where they have overlapping roles ^5,35^. However, in the Met-1 cell line model used in this study we saw no change in Kindlin-2 expression following depletion of Kindlin-1, suggesting that an increase in Kindlin-2 expression is not driving the effects on immune cell populations and anti-tumor immunity. Interestingly Kindlin-2 has been reported to control the recruitment of immunosuppressive (F4/80^+^, CD206^+^) macrophages in orthotopic breast cancer models, where Kindlin-2 in the tumor cells is required for the secretion of colony stimulating factor-1 which acts as a chemoattractant for the macrophages ^36^. Although we did see a reduction in macrophages in the Met-1 tumors lacking Kindlin-1, analysis of phenotypic markers (MHC II, SIRPa and CD206), did not demonstrate any changes in their “polarisation”. Together with our previous work demonstrating the importance of FAK in regulating anti-tumor immunity ^11,37,38^, this highlights the ability of adhesion proteins to control the tumor immune environment. However, Kindlin-1 loss did not mimic the effects of FAK deletion in a model of cutaneous squamous cell carcinoma, indicating context dependency of effects of Kindlin-1 depletion on the anti-tumor immune response. In the skin Kindlin-1 controls epidermal stem cell homeostasis by controlling Wnt and TGF-β signalling leading to a pro-tumorigenic environment ^34^, consistent with the increased susceptibility of Kindler syndrome patients to cutaneous squamous cell carcinoma.

One consistent finding is that the ability of FAK and Kindlin to regulate anti-tumor immunity is dependent on their ability to control cytokine production. Here we show that Kindlin-1 loss leads to increased secretion of IL-6 which reduces the ability of Tregs to suppress CD8+ T cell proliferation. Although IL-6 has well known pro-tumorigenic roles a number of studies have demonstrated the importance of IL-6 in reducing Treg suppressive function in various settings ^28–31^, and also in the differentiation of naïve CD4^+^ T cells into Tregs ^27^. Of note is the observation that IL-6 expression is also increased in keratinocytes from Kindler syndrome patients that lack Kindlin-1 ^9,39^, although its function is not known. The ability of FAK to control expression of the cytokine CCL5 that drives anti-tumor immunity, is mediated via its ability to associate with chromatin where it can complex with transcription factors and their upstream regulators ^11^. Although nuclear roles for Kindlin-2 have been described ^40–42^, there are no reports on potential nuclear functions of Kindlin-1, and it remains to be established how Kindlin-1 controls IL-6, and whether this is conserved in different cell types.

Tregs are suppressive T cells and in addition to their reduced numbers within the tumor microenvironment, we also saw widespread downregulation of numerous functional markers on Tregs in Kindlin-1 depleted tumors that are required for their activation and suppressive activity. Furthermore, an increase in the proportion of resting Tregs (CD26L^+^) compared to activated Tregs (CD62L^-^) in Kin-1 NULL tumors, suggests that Kindlin-1 is also able to modulate their activation state within the tumor microenvironment. These data demonstrate that tumor expression of Kindlin-1 maintains a more suppressive Treg phenotype, which in turn can increase suppression of tumor infiltrating CD8^+^ T cells, with *in vitro* studies identifying a critical role for IL-6 secreted from Kindlin-1 depleted cells in driving the suppressive Treg activity. The presence of Tregs in breast cancer tumors have been associated with poor prognosis and a poor response to immunotherapy in breast cancers ^43,44^. Here we show the importance of Tregs as a method of immune evasion by the Met-1 murine breast cancer model, with depletion of Treg cells (via anti-CD25 antibody) resulting in a reduction in tumor growth, and that this can be controlled by expression of Kindlin-1 in the tumor cells. Further work is needed to assess whether Kindlin-1 is altering the recruitment of Tregs into the tumor, or modulating their differentiation once they have migrated to the tumor microenvironment.

In addition to alterations in Treg function and numbers, an increase in cDC1 cells was also observed in Kindlin-1 depleted tumors. These cells are a subset of DC which express CD103 and are known to be capable of efficient cross-presentation to both CD4 and CD8 cells and have been implicated in generation of T cell driven anti-tumor immune responses ^12,13,45^. Thus, loss of Kindlin-1 impacts on several immune cell types that could contribute to anti-tumor immunity suggesting that it may have multiple roles in controlling immune evasion.

In summary, we have shown that loss of Kindlin-1 results in increased secretion of IL-6, which in turn impedes Treg cell suppressive capacity, resulting in tumor clearance and development of an immunological memory response. These data provide mechanistic insight into how Kindlin-1 expressing tumors can evade immune destruction. As Kindlin-1 is upregulated in breast cancer and linked to survival, targeting Kindlin-1 dependent pathways linked to immune phenotypes may provide a novel strategy to increase the efficacy of immunotherapies in breast cancer, particularly methods relating to reinvigorating the antitumor T cell response.

## Materials and methods

### Cell lines

Met-1 cells were originally acquired from B. Qian (University of Edinburgh) and have been described previously ^46^. Generation of Kin1-WT and Kin1-NULL cells was detailed previously ^7^. Cells were cultured in DMEM high glucose with 10% FBS and hygromycin for selection. Cells were split using TrypLE (Gibco) expression upon 70% confluency. Cells were mycoplasma tested every month and were used within 3 months of recovery from frozen.

### Mice

All experiments were carried out in compliance with UK Home Office regulations. Met-1 Kin1-WT and Kin1-NULL cells (1×10^6^) were injected subcutaneously into both flanks of 8-12 week old female FVB/N mice and tumor growth measured twice weekly using calipers. For tumor growth rechallenge experiments 1×10^6^ cells were injected into FVB/N mice as above. Following tumor regression mice were housed for 35 days prior to rechallenge with 1×10^6^ cells and tumor growth measured. At the time of rechallenge, age matched mice that had not previously been challenged with tumor cells (naïve), were injected with the same cells and tumor growth measured as above. For CD25^+^ cell depletion anti-mouse CD25 depleting antibody (BioXCell, via 2BScientific BP0012) and isotype control (BioXCell, via 2BScientific BP0290) were dosed at 250 μg on day −5, −4 and −3 before 1×10^6^ cells were injected as above on day 0. Antibodies were dosed weekly from day 0 until day 21 at which point the mice were culled and tissue taken for depletion assessment. For anti-PD-L1 blocking, anti-PD-L1 antibody was dosed at 250 μg on day 0, 7, 10, 14 and 17 post 1×10^6^ tumor cell implantation as detailed above.

Generation of MMTV-PyV MMTV-Kin-1^wt/wt^ and MMTV-PyV MMTV-Kin-1^fl/fl^ have been described previously ^7^. Female mice were monitored weekly for tumor formation by palpation.

### Tissue dissociation

After harvesting, tumors were minced and digested with Liberase TL (Roche) and DNase. Tumors were then passed through a 100 μm strainer to achieve a single cell suspension. Spleen and lymph nodes were minced and passed directly through a 100 μm strainer. All tissues underwent red blood cell lysis using RBC lysis buffer (Biolegend).

### Flow cytometry

Following tumor dissociation cells were stained with ZombieUV viability dye (Biolegend), before being resuspended in PBS+1%BSA. Approximately 1×10^6^ cells were aliquoted in 5 ml tubes and incubated with Fc Blocking antibody (Biolegend). Antibodies used for immune profiling are detailed in Supplementary Table 1, and master mixes were prepared before incubation with cells. After washing, cells were fixed and permeabilised overnight using FoxP3/ transcription factor staining buffer set following manufacturer’s instructions (eBiosceince, ThermoFischer), before staining with intracellular antibody master mixes. Finally, cells were washed before being acquired on BD LSR Fortessa (BD Biosciences). Gating and analysis of flow cytometry data was conducted using FlowJo (Version 10.8, Tree Star). Gating of major populations is demonstrated in Supplementary Figure 2 and 4.

### IL-6 ELISA

Met-1 Kin1-WT and Kin1-NULL cells were seeded at 1.5×10^6^ cells per 10 cm^3^ dish. Media was removed after 24 hrs and 5 ml of fresh media was added. After 48 hrs media was harvested, centrifuged and passed through a 0.22 μm filter to remove cell debris. Media was stored at −80 °C until use for maximum of 1 month. If stated, media was concentrated 2X using 3 kDa cut off centrifuge tubes (ThermoFisher) immediately before use. ELISA was conducted using the Mouse IL-6 ELISA Max™ Delux kit (Biolegend), following the manufacturer’s instructions. Plates were read at 450 nm using Tecan Spark 20M plate reader. A reference wavelength reading at 570 nm was subtracted from 450 nm values. Quantification of IL-6 concentration was calculated by extrapolating values using a standard curve of known concentrations.

### Forward phase protein array

Conditioned media was prepared as detailed above. Cells were lysed in RIPA buffer and protein concentration was determined by Pierce BCA protein assay kit (ThermoFischer) following the manufacturer’s instructions. Only proteins which were determined to be above background binding were further analyzed. All values were normalised to protein concentration, and Kin1-NULL values were calculated as fold change over the mean of Kin1-WT values.

### Naïve CD4^+^ differentiation assay

Spleens and lymph nodes (inguinal, axillary, brachial and mesenteric) were harvested from FVB/N mice, minced and passed through a 70 μm strainer. After washing, cells were incubated in RBC lysis buffer to remove red blood cells and resuspended in PBS^+^0.5% BSA^+^EDTA. 1×108 cells were used to isolate by negative selection naïve (CD44^-^) CD4^+^ T cells using Mouse naïve CD4^+^ cell isolation kit (Miltenyi Biotech) following manufactures instructions. 96-well plates were coated overnight in anti-CD3 antibody (7.5 μg/ml; cat# 100340), and anti-CD28 (2 μg/ml; cat#), TGFβ (1 ng/ml; cat#763102) and IL-2 (5 ng/ml, cat# 575402; All Biolegend) were added along with T cell media (RPMI, 10% FBS, 1% L-Glutamine, 0.5% Penicillin-Streptomycin). 5×10^4^ cells per well were added, and stimulated for 5 days in the presence of either Kin1-WT or Kin1-NULL CM (30%). Cells were harvested from plates, added to 5 ml FACS tubes and analyzed by flow cytometry as detailed above.

### Treg suppression assay

Spleens were harvested from FVB/N mice, minced and passed through 70 μm strainer. After washing cells were resuspended in PBS^+^1% BSA and incubated with anti-CD3 (PerCP-Cy5.5; Cat# 100218), anti-CD4 (Brilliant Violet 711™; Cat# 100447), anti-CD8 (Brilliant Violet 510™; Cat# 100100752) and anti-CD25 (PE; Cat# 102008) antibodies. Cells were then sorted using FACS Aria system (BD Biosciences) for CD3^+^ CD8^+^ CD4^-^ CD25^-^ (effector CD8^+^ cells) and CD3^+^ CD8^-^ CD4^+^ CD25^hi^ (Treg cells). Effector CD8^+^ cells were labelled with CellTrace Violet dye (Thermo Fischer) following manufacturer’s instructions. 96-well plates were coated in anti-CD3 antibody (1 μg/ml Biolegend, cat #100340), and splenocytes from CD-1 nude mice were added as antigen presenting cells (APCs). Isolated Tregs and CellTrace violet labelled CD8 cells were co-cultured in T cell media at 1:8 ratio for 5 days in the presence of either Kin1-WT or Kin1-NULL conditioned media (50% concentrated 2X), and with or without anti-IL-6 antibody (20 μg/ml, Biolegend Cat#504512). Cells were then harvested from plates, added to 5 ml FACS tubes and analyzed by flow cytometry as detailed above.

### Nanostring analysis

For bulk RNA analysis: tumors were harvested at day 10 post tumor implantation as described above in ‘Mice’. Tumors were snap frozen in liquid nitrogen and then disrupted and homogenised using RLT buffer (Qiagen). RNA was extracted using RNAeasy kit (Qiagen). For CD45^+^ cell analysis: After harvesting, tumors were processed into a single cell suspension as detailed above in ‘Tissue dissociation’. Cells were incubated with CD45^+^ MACS beads (Miltenyi Biotec) and isolated by positive selection using LS columns (Miltenyi Biotec). 100 μg of RNA was processed using either the mouse Nanostring PanCancer Immune Profiling panel (bulk tumor) or mouse Nanostring Immune Exhaustion Panel (Isolated CD45^+^ cells), following manufacturer’s instructions. Hybridization was performed for 18 hr at 65°C and samples processed using the Nanostring prep station set on high sensitivity. Images were analyzed at maximum (555 fields of view). All data was analyzed by ROSALIND^®^ (https://rosalind.bio/), with a HyperScale architecture developed by ROSALIND, Inc. (San Diego, CA).

### Human data

Pearson correlation of expression of *FERMT1* and *CD274* in all breast cancers the METABRIC microarray dataset (expression log intensity levels), n=1904, r=0.1375 (95% confidence interval 0.09-0.18). Data was downloaded from cbioportal. The figure was made and Pearson correlation and two-tailed t-test were carried out in GraphPad Prism 9.3.0.

### Statistical analysis

Statistical analyses and graphs were produced and performed using a combination of GraphPad Prism version 9.3.0 (GraphPad) and Excel 2016 (Microsoft Corporation). Statistical methods used as detailed in figure legends. Comparisons were considered significantly different when P-value < 0.05.

## Acknowledgements

We would like to thank the Host and Tumor Profiling Unit at the University of Edinburgh Cancer Research UK Centre for help with the Nanostring and forward phase protein arrays. We would also like to thank Marina Jodrell for assistance with performing ELISA assays.

## References

1 Rognoni, E., Ruppert, R. & Fässler, R. The kindlin family: functions, signaling properties and implications for human disease. Journal of cell science 129, 17–27, doi:10.1242/jcs.161190 (2016).

2 Guerrero-Aspizua, S. et al. Assessment of the risk and characterization of non-melanoma skin cancer in Kindler syndrome: study of a series of 91 patients. Orphanet Journal of Rare Diseases 14, 183, doi:10.1186/s13023-019-1158-6 (2019).

3 Lai-Cheong, J. E. et al. Kindler syndrome: a focal adhesion genodermatosis. The British journal of dermatology 160, 233–242, doi:10.1111/j.1365-2133.2008.08976.x (2009).

4 Zhan, J. & Zhang, H. Kindlins: Roles in development and cancer progression. The International Journal of Biochemistry & Cell Biology 98, 93–103, doi:https://doi.org/10.1016/j.biocel.2018.03.008 (2018).

5 Azorin, P. et al. Distinct expression profiles and functions of Kindlins in breast cancer. Journal of Experimental & Clinical Cancer Research 37, 281, doi:10.1186/s13046-018-0955-4 (2018).

6 Sin, S.et al. Role of the Focal Adhesion Protein Kindlin-1 in Breast Cancer Growth and Lung Metastasis. JNCI: Journal of the National Cancer Institute 103, 1323–1337, doi:10.1093/jnci/djr290 (2011).

7 Sarvi, S. et al. Kindlin-1 Promotes Pulmonary Breast Cancer Metastasis. Cancer research 78, 1484–1496, doi:10.1158/0008-5472.can-17-1518 (2018).

8 Heinemann, A. et al. Induction of phenotype modifying cytokines by FERMT1 mutations. Human Mutation 32, 397–406, doi:https://doi.org/10.1002/humu.21449 (2011).

9 Maier, K. et al. UV-B-induced cutaneous inflammation and prospects for antioxidant treatment in Kindler syndrome. Human Molecular Genetics 25, 5339–5352, doi:10.1093/hmg/ddw350 (2016).

10 Chacón-Solano, E. et al. Fibroblast activation and abnormal extracellular matrix remodelling as common hallmarks in three cancer-prone genodermatoses. The British journal of dermatology 181, 512–522, doi:10.1111/bjd.17698 (2019).

11 Serrels, A. et al. Nuclear FAK controls chemokine transcription, Tregs, and evasion of anti-tumor immunity. Cell 163, 160–173, doi:10.1016/j.cell.2015.09.001 (2015).

12 Ferris, S. T. et al. cDC1 prime and are licensed by CD4(+) T cells to induce antitumour immunity. Nature 584, 624–629, doi:10.1038/s41586-020-2611-3 (2020).

13 Laoui, D. et al. The tumour microenvironment harbours ontogenically distinct dendritic cell populations with opposing effects on tumour immunity. Nature communications 7, 13720, doi:10.1038/ncomms13720 (2016).

14 Zhou, F. Molecular mechanisms of IFN-gamma to up-regulate MHC class I antigen processing and presentation. International reviews of immunology 28, 239–260, doi:10.1080/08830180902978120 (2009).

15 Garcia-Diaz, A. et al. Interferon Receptor Signaling Pathways Regulating PD-L1 and PD-L2 Expression. Cell Rep 19, 1189–1201, doi:10.1016/j.celrep.2017.04.031 (2017).

16 Hatzioannou, A. et al. Regulatory T Cells in Autoimmunity and Cancer: A Duplicitous Lifestyle. Frontiers in immunology 12, 731947, doi:10.3389/fimmu.2021.731947 (2021).

17 Luo, C. T., Liao, W., Dadi, S., Toure, A. & Li, M. O. Graded Foxo1 activity in Treg cells differentiates tumour immunity from spontaneous autoimmunity. Nature 529, 532–536, doi:10.1038/nature16486 (2016).

18 Ren, J. et al. Foxp1 is critical for the maintenance of regulatory T-cell homeostasis and suppressive function. PLoS Biol 17, e3000270 (2019). <http://europepmc.org/abstract/MED/31125332>.

19 Huehn, J. et al. Developmental Stage, Phenotype, and Migration Distinguish Naive- and Effector/Memory-like CD4+ Regulatory T Cells. Journal of Experimental Medicine 199, 303–313, doi:10.1084/jem.20031562 (2004).

20 Willoughby, J., Griffiths, J., Tews, I. & Cragg, M. S. OX40: Structure and function - What questions remain? Molecular immunology 83, 13–22, doi:10.1016/j.molimm.2017.01.006 (2017).

21 Alissafi, T., Hatzioannou, A., Legaki, A. I., Varveri, A. & Verginis, P. Balancing cancer immunotherapy and immune-related adverse events: The emerging role of regulatory T cells. Journal of Autoimmunity 104, 102310, doi:https://doi.org/10.1016/j.jaut.2019.102310 (2019).

22 Antonioli, L., Pacher, P., Vizi, E. S. & Haskó, G. CD39 and CD73 in immunity and inflammation. Trends in Molecular Medicine 19, 355–367, doi:https://doi.org/10.1016/j.molmed.2013.03.005 (2013).

23 Allard, B., Longhi, M. S., Robson, S. C. & Stagg, J. The ectonucleotidases CD39 and CD73: Novel checkpoint inhibitor targets. Immunological reviews 276, 121–144, doi:10.1111/imr.12528 (2017).

24 Hirano, N. et al. Engagement of CD83 ligand induces prolonged expansion of CD8+ T cells and preferential enrichment for antigen specificity. Blood 107, 1528–1536, doi:10.1182/blood-2005-05-2073 (2006).

25 Nicolet, B. P. et al. CD29 identifies IFN-γ-producing human CD8(+) T cells with an increased cytotoxic potential. Proceedings of the National Academy of Sciences of the United States of America 117, 6686–6696, doi:10.1073/pnas.1913940117 (2020).

26 Kazanietz, M. G., Durando, M. & Cooke, M. CXCL13 and Its Receptor CXCR5 in Cancer: Inflammation, Immune Response, and Beyond. Frontiers in endocrinology 10, 471, doi:10.3389/fendo.2019.00471 (2019).

27 Kimura, A. & Kishimoto, T. IL-6: regulator of Treg/Th17 balance. European journal of immunology 40, 1830–1835, doi:10.1002/eji.201040391 (2010).

28 Guo, H. et al. Stability and inhibitory function of Treg cells under inflammatory conditions in vitro. Experimental and therapeutic medicine 18, 2443–2450, doi:10.3892/etm.2019.7873 (2019).

29 Yang, X. O. et al. Molecular antagonism and plasticity of regulatory and inflammatory T cell programs. Immunity 29, 44–56, doi:10.1016/j.immuni.2008.05.007 (2008).

30 Ye, M. et al. Deletion of IL-6 Exacerbates Colitis and Induces Systemic Inflammation in IL-10-Deficient Mice. Journal of Crohn’s & colitis 14, 831–840, doi:10.1093/ecco-jcc/jjz176 (2020).

31 Garg, G. et al. Blimp1 Prevents Methylation of <em>Foxp3</em> and Loss of Regulatory T Cell Identity at Sites of Inflammation. Cell Reports 26, 1854–1868.e1855, doi:10.1016/j.celrep.2019.01.070 (2019).

32 Ruan, Q. et al. The Th17 immune response is controlled by the Rel-RORγ-RORγ T transcriptional axis. The Journal of experimental medicine 208, 2321–2333, doi:10.1084/jem.20110462 (2011).

33 Patel, H. et al. Kindlin-1 regulates mitotic spindle formation by interacting with integrins and Plk-1. Nature communications 4, 2056, doi:10.1038/ncomms3056 (2013).

34 Rognoni, E. et al. Kindlin-1 controls Wnt and TGF-β availability to regulate cutaneous stem cell proliferation. Nature Medicine 20, 350–359, doi:10.1038/nm.3490 (2014).

35 He, Y., Esser, P., Heinemann, A., Bruckner-Tuderman, L. & Has, C. Kindlin-1 and −2 have overlapping functions in epithelial cells implications for phenotype modification. The American journal of pathology 178, 975–982, doi:10.1016/j.ajpath.2010.11.053 (2011).

36 Sossey-Alaoui, K. et al. Kindlin-2 Regulates the Growth of Breast Cancer Tumors by Activating CSF-1-Mediated Macrophage Infiltration. Cancer research 77, 5129–5141, doi:10.1158/0008-5472.can-16-2337 (2017).

37 Canel, M. et al. T-cell co-stimulation in combination with targeting FAK drives enhanced anti-tumor immunity. eLife 9, e48092, doi:10.7554/eLife.48092 (2020).

38 Serrels, B. et al. IL-33 and ST2 mediate FAK-dependent antitumor immune evasion through transcriptional networks. Science Signaling 10, eaan8355, doi:doi:10.1126/scisignal.aan8355 (2017).

39 Qu, H., Wen, T., Pesch, M. & Aumailley, M. Partial loss of epithelial phenotype in kindlin-1-deficient keratinocytes. The American journal of pathology 180, 1581–1592, doi:10.1016/j.ajpath.2012.01.005 (2012).

40 Yu, Y. et al. Kindlin 2 promotes breast cancer invasion via epigenetic silencing of the microRNA200 gene family. International Journal of Cancer 133, 1368–1379, doi:https://doi.org/10.1002/ijc.28151 (2013).

41 Wang, P. et al. Kindlin-2 interacts with and stabilizes DNMT1 to promote breast cancer development. The International Journal of Biochemistry & Cell Biology 105, 41–51, doi:https://doi.org/10.1016/j.biocel.2018.09.022 (2018).

42 Sossey-Alaoui, K., Pluskota, E., Szpak, D. & Plow, E. F. The Kindlin2-p53-SerpinB2 signaling axis is required for cellular senescence in breast cancer. Cell Death & Disease 10, 539, doi:10.1038/s41419-019-1774-z (2019).

43 Núñez, N. G. et al. Tumor invasion in draining lymph nodes is associated with Treg accumulation in breast cancer patients. Nature communications 11, 3272, doi:10.1038/s41467-020-17046-2 (2020).

44 Plitas, G. et al. Regulatory T Cells Exhibit Distinct Features in Human Breast Cancer. Immunity 45, 1122–1134, doi:10.1016/j.immuni.2016.10.032 (2016).

45 Noubade, R., Majri-Morrison, S. & Tarbell, K. V. Beyond cDC1: Emerging Roles of DC Crosstalk in Cancer Immunity. Frontiers in immunology 10, 1014, doi:10.3389/fimmu.2019.01014 (2019).

46 Borowsky, A. D. et al. Syngeneic mouse mammary carcinoma cell lines: two closely related cell lines with divergent metastatic behavior. Clinical & experimental metastasis 22, 47–59, doi:10.1007/s10585-005-2908-5 (2005).

