## Supplementary Tables and figures for "Loss of the adhesion protein Kindlin-1 stimulates tumor clearance via modulation of Tregs"

| Antigen | Flurophore | Supplier | Cat # |
|---|---|---|---|
| CD3 | PerCP-Cy5.5 | Biolegend | 100218 |
| CD3 | BUV496 | BD Biosciences | 612955 |
| CD4 | Brilliant Violet 711™ | Biolegend | 100447 |
| CD4 | APC-H7 | BD Biosciences | 560246 |
| CD8 | Brilliant Violet 510™ | Biolegend | 100752 |
| FoxP3 | PE | ThermoFischer | 12-5773-82 |
| γδTCR | Brilliant Violet 421™ | Biolegend | 118120 |
| CD44 | APC-R700 | BD Biosciences | 565480 |
| CD62L | PE-Cy7 | Biolegend | 104418 |
| RoRγT | BV650 | BD Biosciences | 564722 |
| CD25 | Brilliant Violet 785™ | Biolegend | 102051 |
| PD-1 | Brilliant Violet 605™ | Biolegend | 135220 |
| TIM-3 | APC | ThermoFischer | 17-5871-80 |
| CD45 | BV786 | BD Biosciences | 564225 |
| CD11b | APC-R700 | BD Biosciences | 564985 |
| CD11c | PE/Dazzle™ 594 | Biolegend | 117348 |
| F4/80 | FITC | Biolegend | 123108 |
| Ly6C | PerCP-Cy5.5 | Biolegend | 128012 |
| Ly6G | PE-Cy7 | Biolegend | 108415 |
| MHC II | Brilliant Violet 510™ | Biolegend | 107636 |
| CD206 | Brilliant Violet 711™ | Biolegend | 141727 |
| PD-L1 | Brilliant Violet 650™ | Biolegend | 124336 |
| CD103 | Brilliant Violet 605™ | Biolegend | 121433 |
| SIRPa | PE | Biolegend | 144012 |
| CLEC9a | BV421 | BD Biosciences | 564271 |
| GITR | FITC | Biolegend | 120205 |
| TIGIT | Brilliant Violet 421™ | Biolegend | 142111 |
| OX40 | PerCP-Cy5.5 | Biolegend | 119424 |
| CD83 | Brilliant Violet 650™ | Biolegend | 121515 |
| 4-1BB | eFluor450 | ThermoFischer | 48-1371-80 |
| CTLA-4 | PerCP-Cy5.5 | Biolegend | 106315 |
| LAG3 | BV711 | BD Biosciences | 563179 |
| CD39 | APC | Biolegend | 143809 |
| CD73 | Brilliant Violet 605™ | Biolegend | 127215 |
| CD29 | APC-Cy7 | Biolegend | 102225 |

**Supplementary Table 1 – List of anti-mouse antibodies used for flow cytometry**

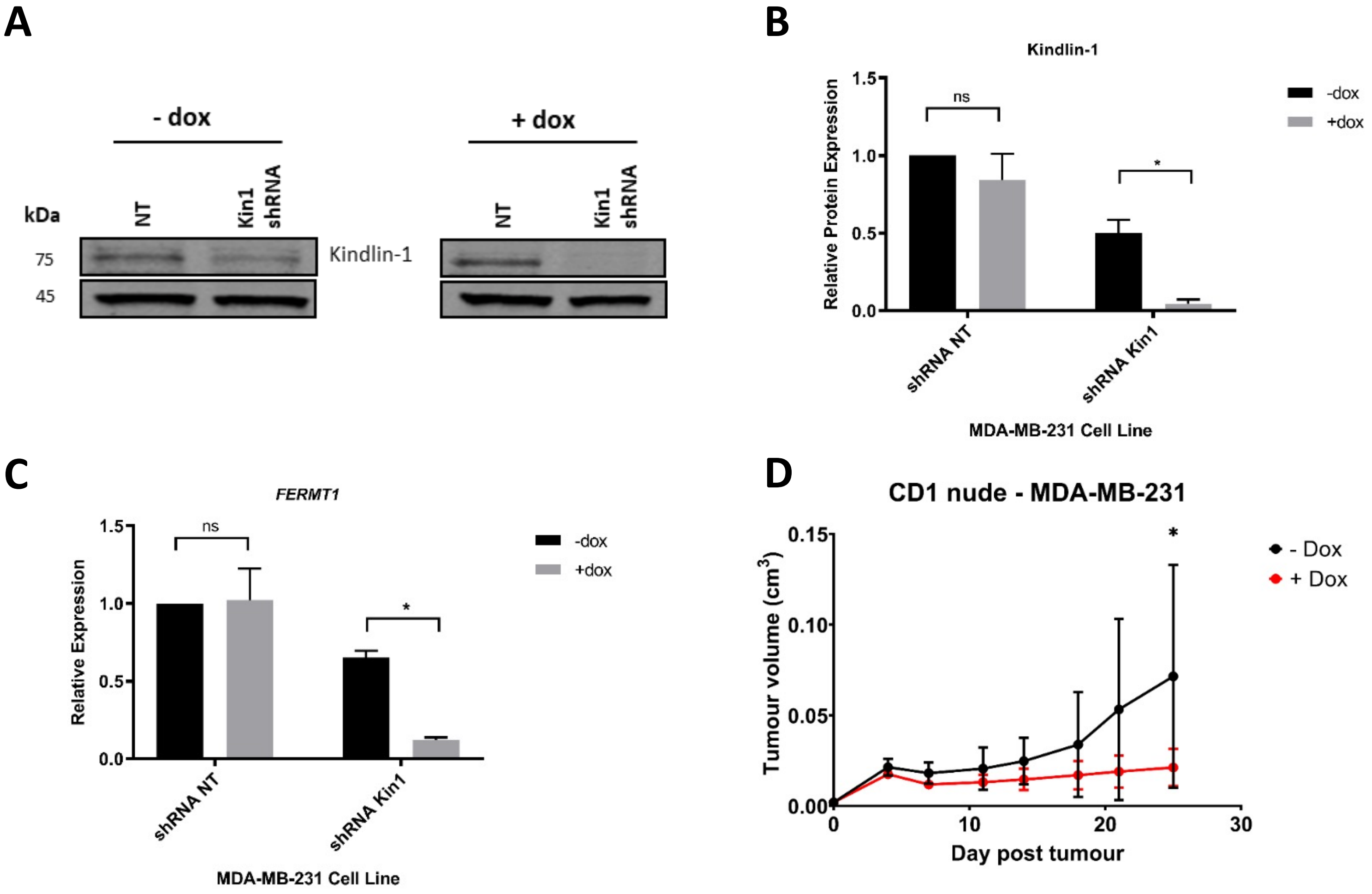


**Supplementary Figure 1 – Loss of Kindlin-1 leads to reduction of tumor growth in human breast cancer model A)** Representative western blot demonstrating Kindlin-1 protein knockdown after addition of doxycycline to MDA-MB-231 cells expressing inducible *FERMT1* shRNA. **B)** Quantification of A normalised to loading control. **C)** qPCR quantification of *FERMT1* (Kindlin-1) after shRNA knockdown in MDA-MB-231 cells. **D)** MDA-MB-231 cells carrying doxycycline inducible *FERMT1* shRNA were subcutaneously injected into CD1 nude mice, after which doxycycline treatment was commenced for half the group. Tumor growth was monitored and recorded until day 25, with average tumor growth shown. Unpaired t-test with *= <0.05, ** = <0.01, *** = <0.001.

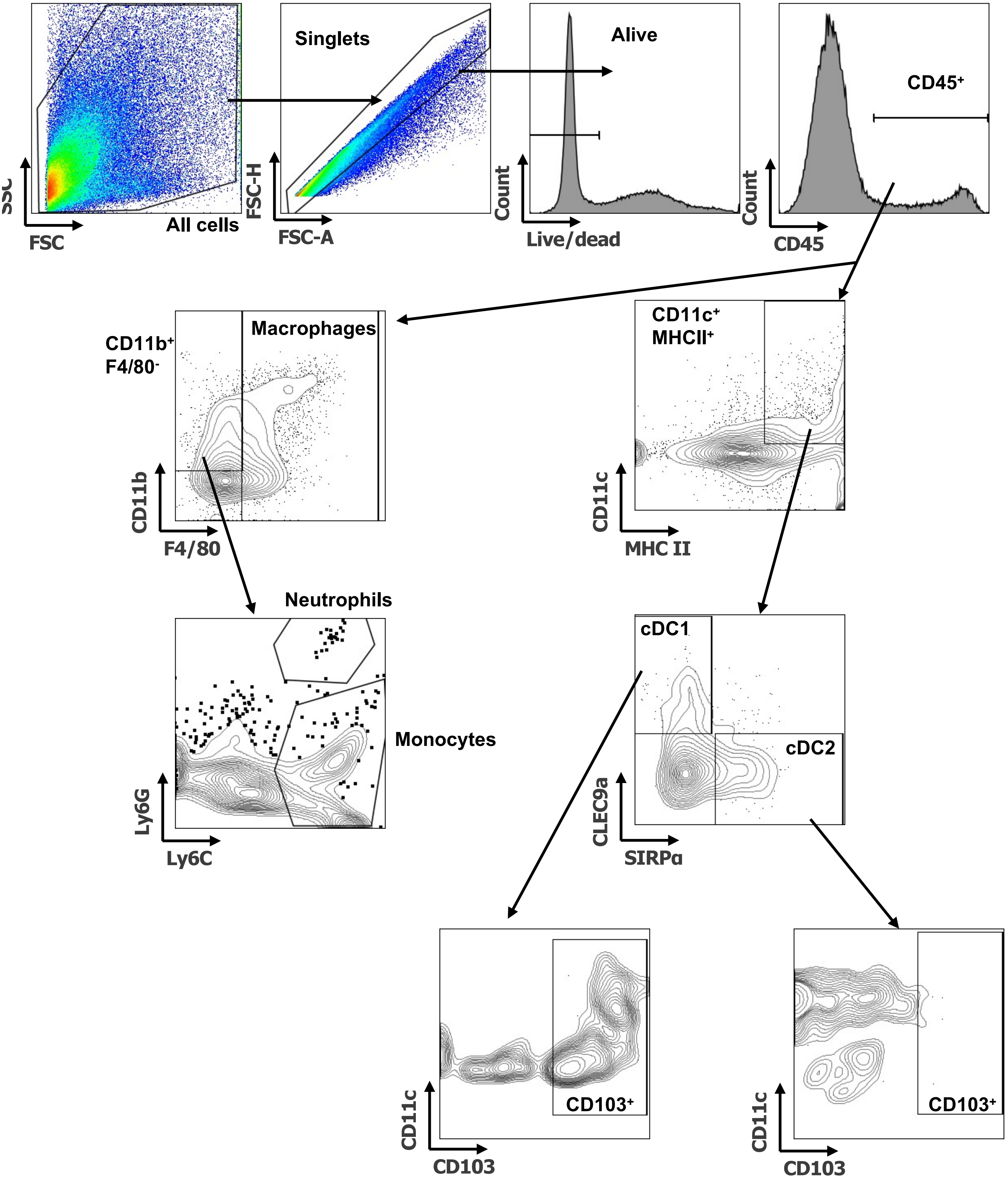


**Supplementary Figure 2 – Flow cytometry gating examples of myeloid populations.** Example of tumor myeloid cell gating shown. First debris is removed by All cells gate, followed by singlets and live cells. CD45+ are selected for downstream identification of Macrophages (F4/80+), Neutrophils (CD11b+ F4/80- Ly6Ghi Ly6cint) and Monocytes (CD11b+ F4/80- Ly6Glo Ly6C+). Total dendritic cells (DCs) were defined as CD11c+ MHC II+ with further downstream gating of cDC1 (CLEC9a+ SIRPα-) and cDC2 (CLEC9a- SIRPα+). Expression of CD103 was then assessed on cDC1s and cDC2s.

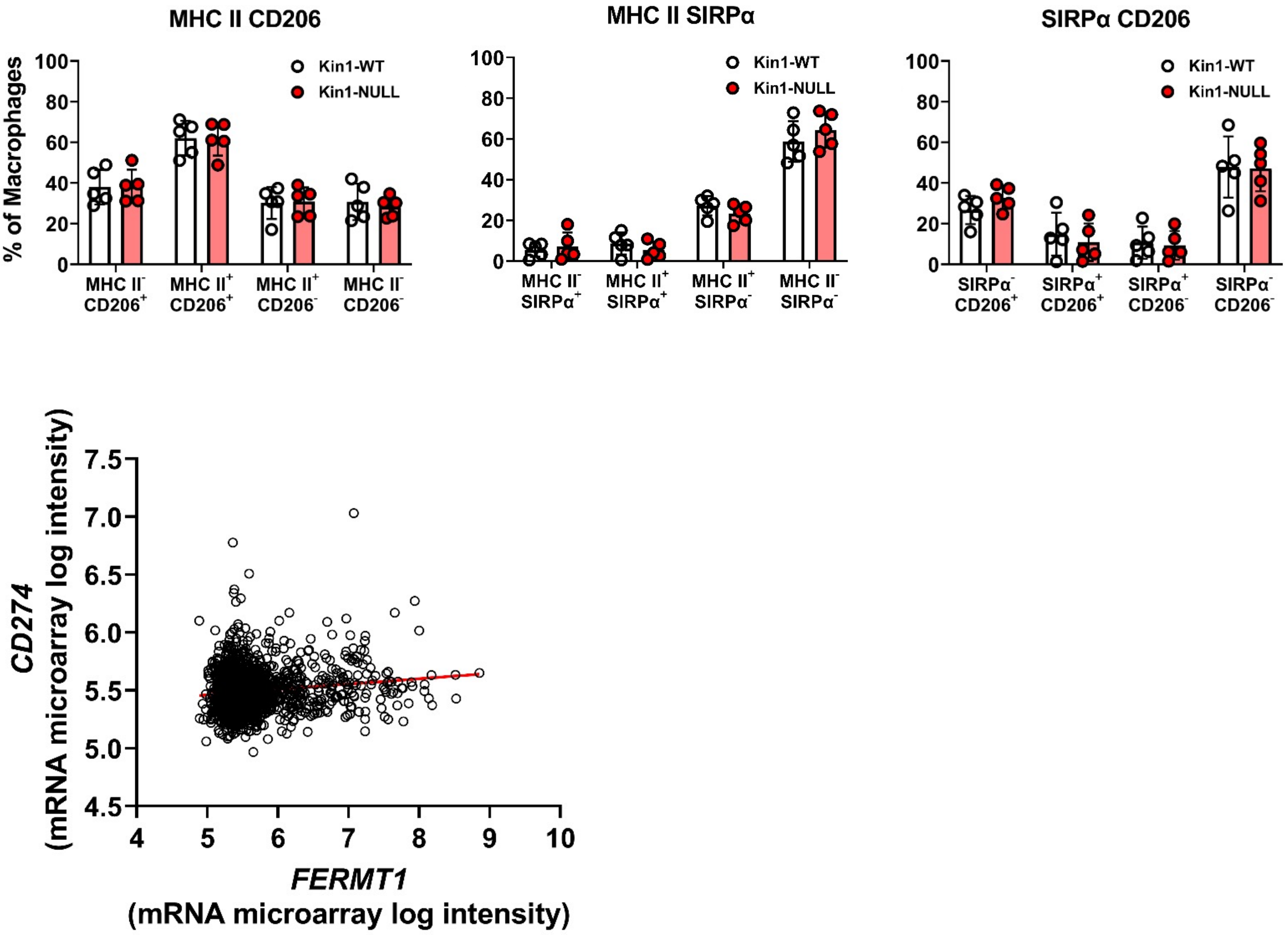


**Supplementary Figure 3 – Macrophage phenotype profiling in Met-1 tumors and PD-L1 Kindlin-1 correlation in human breast cancer dataset. A)** Quantification of MHC II and CD206 (left), MHC II and SIRPα (middle), and SIRPα and CD206 (right) as a percentage of total macrophages infiltrating MET-01 tumours. Data representative of two independent experiments. n=5 per group. **B)** Correlation analysis of *FERMT1* (Kindlin-1) and *CD274* (PD-L1) using METABRIC human breast cancer data set. n=1904, r=0.1375 (95% confidence interval 0.09-0.18), p<0.0001.

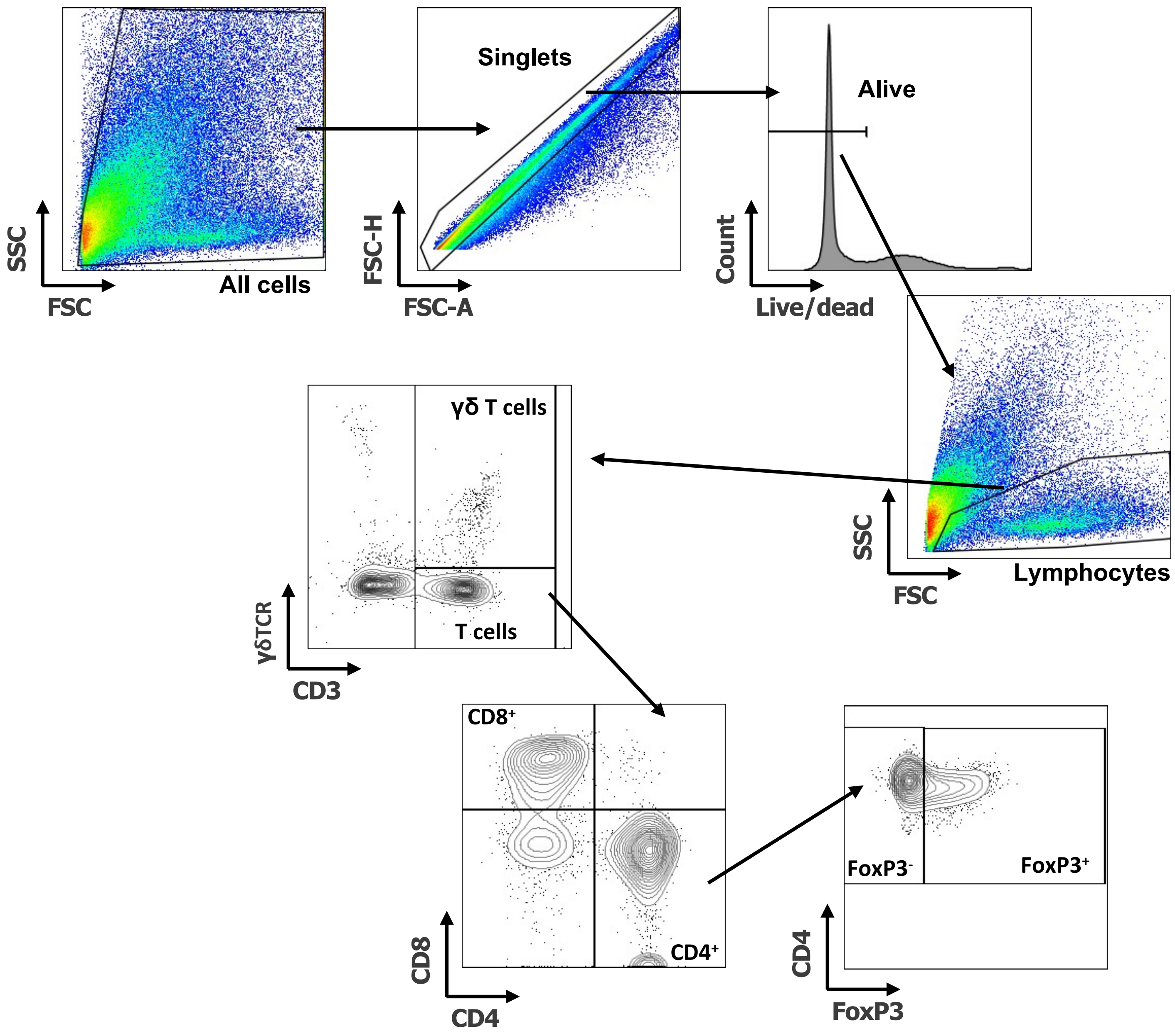


**Supplementary Figure 4 – Flow cytometry gating examples of T cell populations.** Example of tumor T cell cell gating shown. First debris is removed by All cells gate, followed by singlets, live cells and lymphocytes. γδT cells (CD3+ γδTCR+) and T cells (CD3+ γδTCR-) were gated. T cells were then further subdivided into CD8+ and CD4+ T cells. From the CD4+ gate Tregs (FoxP3+) and non-Treg CD4s (FoxP3-) were gated.

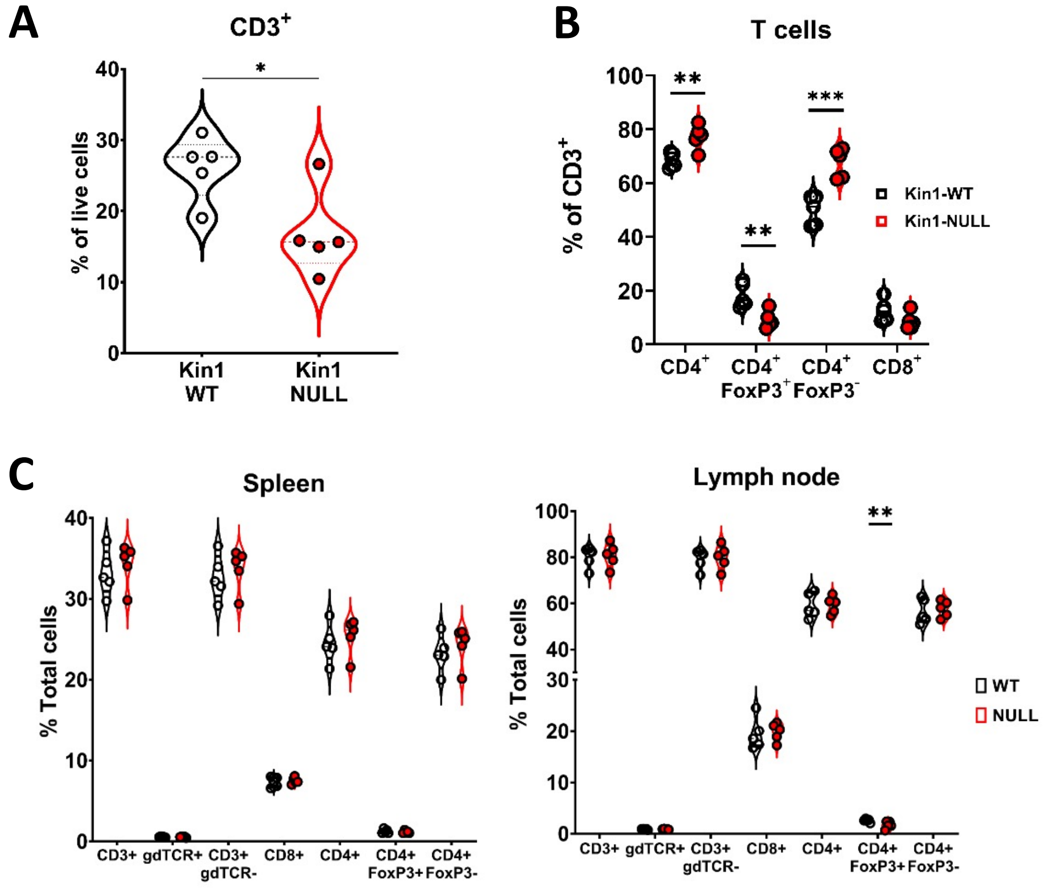


**Supplementary Figure 5 – Loss of Kindlin-1 reduces tumor infiltrating Treg cells. A)** Met-1 Kin1-WT or Kin1-NULL tumors were established via subcutaneous injection in FVB mice, and harvested at day 10 for immunophenotyping by flow cytometry. Gating of CD3+ cells was conducted with subsequent gating and quantification of major T cell populations (**B**) as percentage of total CD3+ cells. **C)** As in A but quantification of T cell subset in spleen (left) and draining (inguinal) lymph nodes (right). Example of two independent experiments (A-C). n=3-5 per group. Unpaired t-test with *= <0.05, ** = <0.01, *** = <0.001.

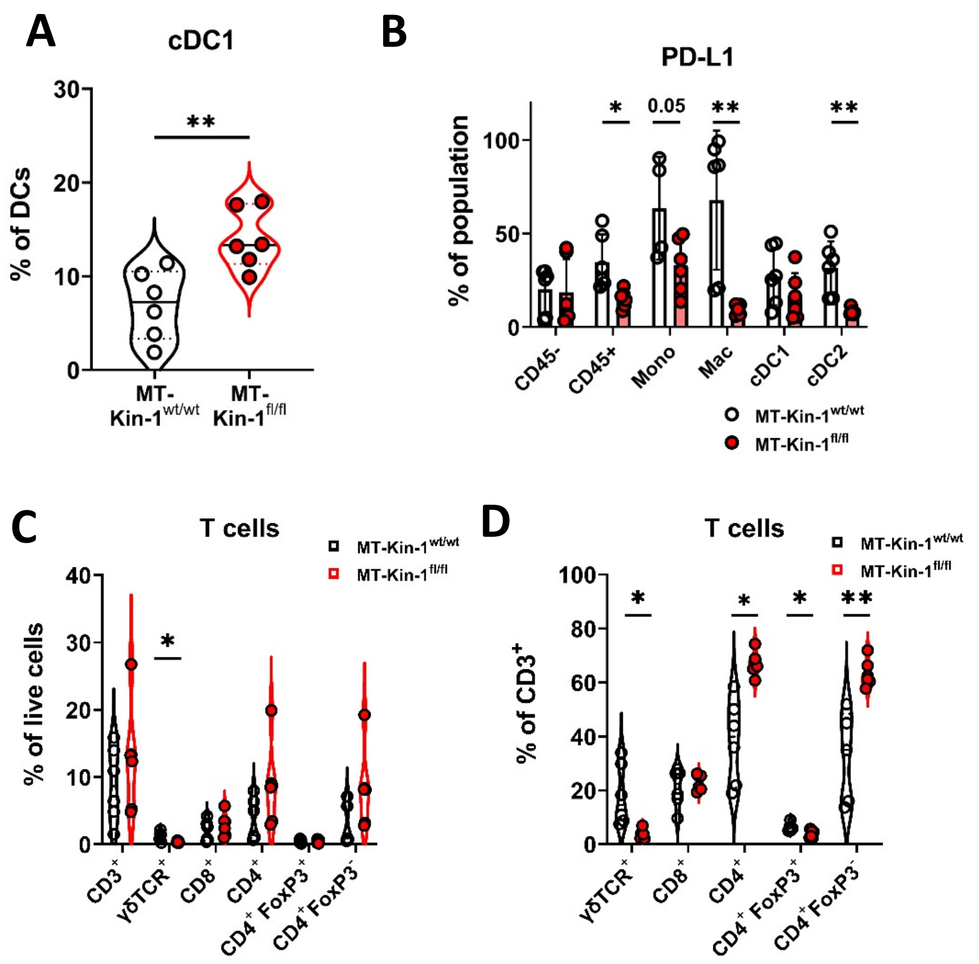


**Supplementary Figure 6 – Immune modulation in MMTV-PyV MMTV-Kin-1$^{wt/wt}$ and MMTV-Kin-1$^{fl/fl}$ spontaneous tumor model. A)** Spontaneous MT-Kin-1$^{wt/wt}$ or MT-Kin-1$^{fl/fl}$ tumors were harvested once 1 tumour reached 10 mm for immunophenotyping by flow cytometry. Quantification of cDC1 cells as a percentage of total DCs (CD11c+ MHC II+). **B)** Quantification of PD-L1 expression on non-immune (CD45$^{-}$) and myeloid subsets from MT-Kin-1$^{wt/wt}$ or MT-Kin-1$^{fl/fl}$ tumours, as a percentage of the corresponding cell population. **C)** As in A but quantification of major T cell subsets as a % of live (total) cells and **D)** CD3+ cells. n=5-6 mice per group. Unpaired t-test with *= <0.05, ** = <0.01.

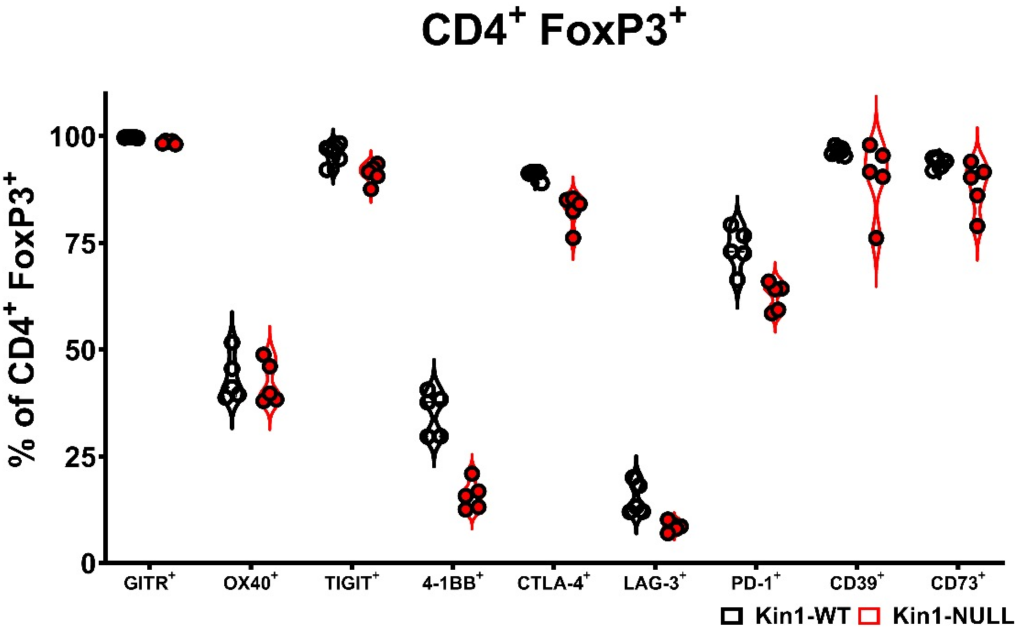


**Supplementary Figure 7 – Loss of Kindlin-1 modulates Treg phenotype markers.** Met-1 Kin1-WT or Kin1-NULL tumors were established via subcutaneous injection in FVB mice, and harvested at day 10 for immunophenotyping by flow cytometry. Analysis of expression of markers were assessed on Tregs (CD4+ FoxP3+). Expression shown as a percentage of population. n=4-5 per group.

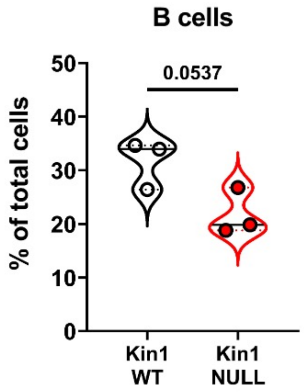


**Supplementary Figure 8 – Quantification of B cells in Met-1 Kin1-WT and Kin1-NULL tumors.** Met-1 Kin1-WT or Kin1-NULL tumors were established via subcutaneous injection in FVB mice, and harvested at day 10 for immunophenotyping by flow cytometry. Gating of $CD3^-$ cells was conducted with subsequent gating of B220+ B cells as a percentage of total cells. n=3 per group. Unpaired t-test.

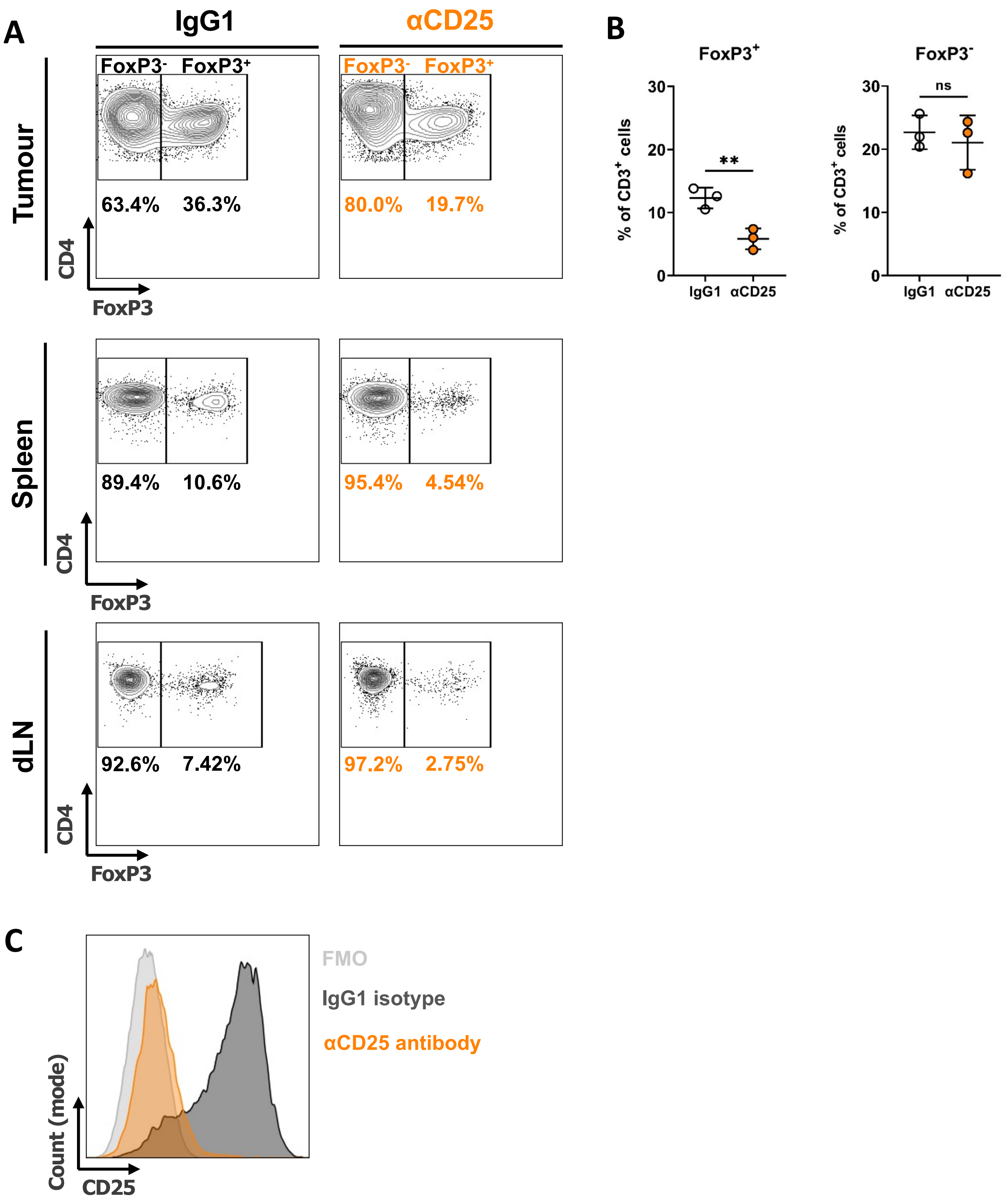


**Supplementary Figure 9 – Depletion of Treg with anti-CD25 antibody treatment. A)** After 3 day pre treatment with anti-CD25 or isotype control antibody, Met-1 Kin1-WT or Kin1-NULL tumors were established via subcutaneous injection in FVB mice. Antibody treatment was continued weekly for 21 days at which point tissue was harvested to assess Treg specific deletion. Representative plots showing depletion of Treg cells in tumour (top), spleen (middle) and draining lymph node (dLN – bottom). **B)** Quantification of tumour data for both FoxP3+ (Treg) and FoxP3- (non-Treg CD4+) cells. **C)** Example histogram demonstrating CD25 expression on Treg cells. Representative example of two experiments. n=3 per group. Unpaired t-test *= <0.05, ** = <0.01.
